# The dendritic spatial code: branch-specific place tuning and its experience-dependent decoupling

**DOI:** 10.1101/2020.01.24.916643

**Authors:** Shannon K. Rashid, Victor Pedrosa, Martial A. Dufour, Jason J. Moore, Spyridon Chavlis, Rodrigo G. Delatorre, Panayiota Poirazi, Claudia Clopath, Jayeeta Basu

**Affiliations:** New York University Neuroscience Institute, New York University, New York, NY 10016, USA; Department of Neuroscience and Physiology, New York University School of Medicine, New York, NY 10016, USA; Department of Bioengineering, Imperial College London, London, SW7 2AZ, UK; CAPES Foundation, Ministry of Education of Brazil, Brasilia, 70040-020, Brazil; Institute of Molecular Biology and Biotechnology (IMBB), Foundation for Research and Technology-Hellas (FORTH), Heraklion, Crete, Greece; Center for Neural Science, New York University, New York, NY 10003, USA; Department of Psychiatry, New York University School of Medicine, New York, NY 10016, USA

## Abstract

Dendrites of pyramidal neurons integrate different sensory inputs, and non-linear dendritic computations drive feature selective tuning and plasticity. Yet little is known about how dendrites themselves represent the environment, the degree to which they are coupled to their soma, and how that coupling is sculpted with experience. In order to answer these questions, we developed a novel preparation in which we image soma and connected dendrites in a single plane across days using *in vivo* two-photon microscopy. Using this preparation, we monitored spatially tuned activity in area CA3 of the hippocampus in head-fixed mice running on a linear track. We identified “place dendrites”, which can stably and precisely represent both familiar and novel spatial environments. Dendrites could display place tuning independent of their connected soma and even their sister dendritic branches, the first evidence for branch-specific tuning in the hippocampus. In a familiar environment, spatially tuned somata were more decoupled from their dendrites as compared to non-tuned somata. This relationship was absent in a novel environment, suggesting an experience dependent selective gating of dendritic spatial inputs. We then built a data-driven multicompartment computational model that could capture the experimentally observed correlations. Our model predicts that place cells exhibiting branch-specific tuning have more flexible place fields, while neurons with homogenous or co-tuned dendritic branches have higher place field stability. These findings demonstrate that spatial representation is organized in a branch-specific manner within dendrites of hippocampal pyramidal cells. Further, spatial inputs from dendrites to soma are selectively and dynamically gated in an experience-dependent manner, endowing both flexibility and stability to the cognitive map of space.

**One sentence summary:** Hippocampal pyramidal cells show branch-specific tuning for different place fields, and their coupling to their soma changes with experience of an environment.

## Results

Pyramidal neurons across the brain integrate inputs from multiple sources upon their dendritic arbors. However, it is widely observed that *somatic* activity of these neurons is highly selective for features of the external environment. How can a neuron with variable input exhibit selective output? Despite years of research regarding the organization and feature selectivity of these neural representations at the somatic level, the subcellular computations underlying them remains largely elusive. Although somata tend to be tuned to only one feature, at least two studies in the cortex show branch-specific tuning, whereby sensory (*1*) and motor (*2*) features are differentially represented even by sister dendritic branches. Does the soma integrate these varied inputs from different dendrites, or does it select for certain inputs over others to dictate the executive output of the neuron?

A prominent feature selectivity observed in the hippocampus across many mammalian species is the spatial tuning of place cells (*3, 4*). Place cells show highly selective activity at distinct locations of space (place fields) as an animal navigates its environment (*5*). These neurons derive spatial information from both the entorhinal cortex and within the hippocampus through inputs upon their dendrites. These inputs are compartmentalized such that different input streams target different segments of the dendritic tree (*6-10*). Although we lack experimental evidence for dendritic branch-specific tuning in hippocampal neurons, place cells provide an ideal substrate to study the relationship between dendritic and somatic signals. If individual dendritic branches of a given place cell were selectively tuned to distinct spatial locations, it would allow a single place cell to hold multiple pieces of spatial information. This would vastly increase the computational power of the single cell, enabling the rapid formation of tuned place cells as an animal explores a novel environment (*11-14*) or the remapping of existing place fields to report distinctions between environments (*12-15*).

Non-linear activity such as dendritic spikes enhance feature selectivity in the visual cortex (*16*) and drive long term synaptic plasticity even independent of somatic spiking in the barrel cortex (*17*). Non-linear dendritic spikes have long been implicated in the induction of cooperative long-term plasticity *in vitro* (*18-21*) and complex spike burst activity in the soma *in vivo* (*22-24*), a signature feature of place cells and synaptic plasticity. In area CA1 of the hippocampus, intracellular somatic recordings reveal that injected subthreshold depolarization as well as dendritic spike-like waveforms (*22*) are capable of converting silent cells into place cells. Further, multi-planar imaging of CA1 pyramidal neurons and basal dendrites shows that greater prevalence of dendritic spikes across multiple branches correlates with better place field precision and stability (*26*). Despite these pathbreaking findings emphasizing the role of dendritic spikes in predicting the spatial tuning of place cells, there remains a fundamental gap in our understanding of the dendrite to soma transformation relationship and the subcellular organization of spatial representation in the long-term. This gap stems from a lack of experimental data that can directly and simultaneously read out the activity of place cell somata and large expanses of their connected apical and basal dendrites over extended periods of time. Such experimental data would enable us to unequivocally address important unanswered questions: How compartmentalized is the representation of the spatial environment at the level of a single cell? Do individual dendritic branches represent place information in a long-term stable manner? Are they capable of having independent place fields from their soma? As in the cortex (*1, 2*), can hippocampal place cells show branch-specific tuning to spatial locations? And if so, how do place cells that typically show single place fields in their somatic output, integrate and/or gate variable inputs from their dendrites?

To fill this gap we developed a cranial window preparation (Fig 1A-B, D), that captures a novel field of view with somata and connected basal and apical dendrites of the same cells in a single plane: allowing us to perform multi-compartment imaging at high speed (29 fps) and resolution (1.085 µm/ pixel) in head-fixed mice running on a linear treadmill (*27-29*)(Fig 1C).

**Figure 1:**
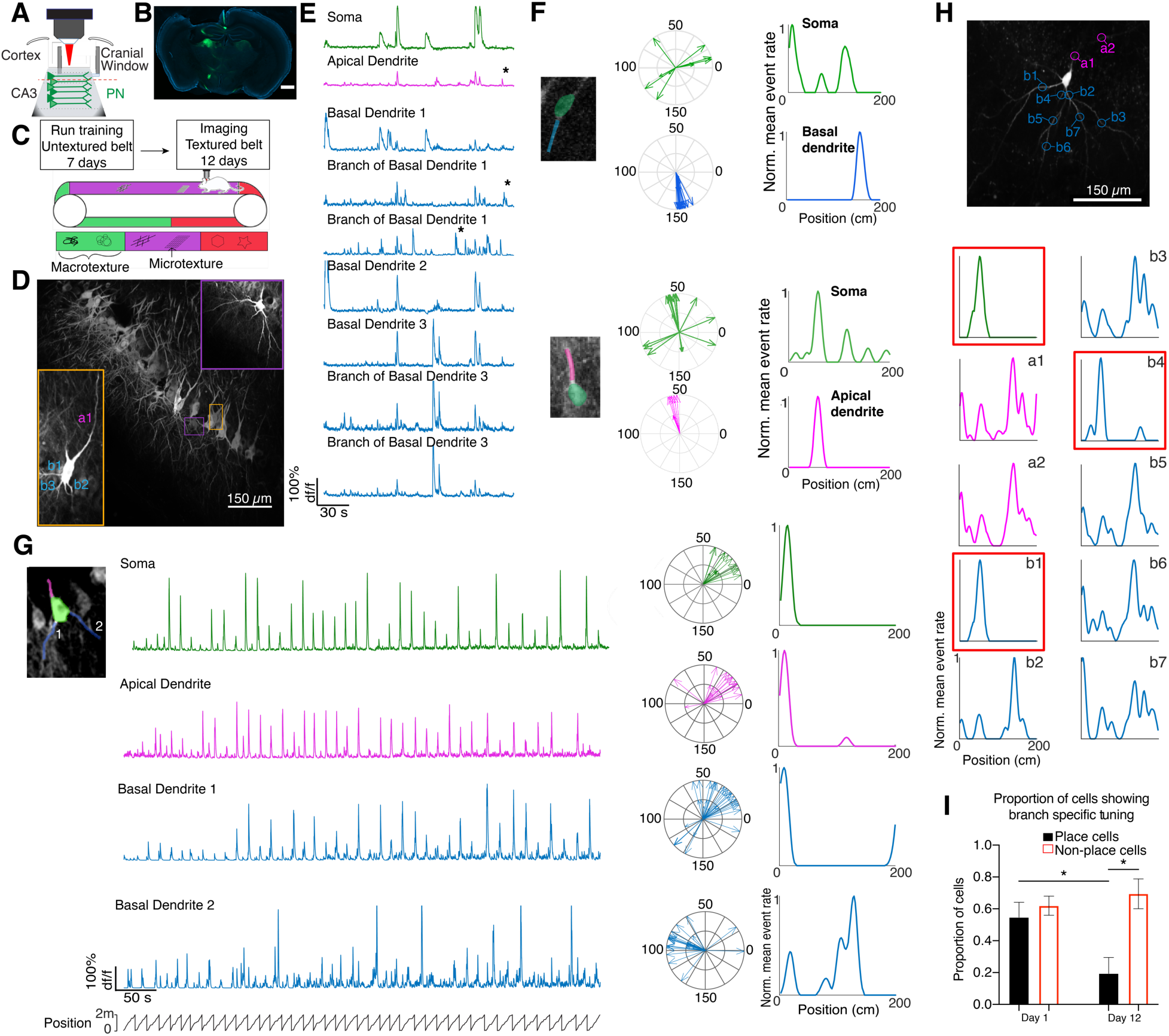
Imaging preparation reveals branch-specific tuning. **A.** Schematic of CA3 cranial window. **B.** Coronal slice from an animal that underwent an injection of GCaMP6f into CA3 and cranial window implantation. Scale bar = 400 µm. **C.** Schematic of experimental paradigm. **D.** Example field of view, insets showing dendritic branching of two example cells. **E.** Example traces from cell boxed in orange in D. Green traces are somatic, magenta are apical dendrites, and blue are basal dendrites. Asterisks denote examples of events generated in the dendrite. **F.** Example of independently tuned basal dendrite (top) and an example independently tuned apical dendrite (bottom). Each arrow represents a significant calcium event for the session shown. **G.** Example of tuned dendrites from a cell that exhibits branch-specific tuning. The soma and each dendrite pictured all meet spatial tuning criteria. Basal dendrite 2 has a different place field than the others. **H.** Example of branch-specific tuning for a cell from an animal that had sparse GCaMP6f expression, ensuring non-overlapping labeled dendrites. **I.** Proportion of tuned and non-tuned soma showing branch-specific tuning on day 1 and day 12 of imaging (* denotes a p < 0.05)

To monitor calcium activity of both somata and dendrites (Fig 1E), we virally expressed GCaMP6f in pyramidal neurons of CA3 (Fig S1), an area that harbors place cell ensembles (*5*) exhibiting high degrees of stability as well as flexibility in a context-dependent manner (*14, 30*). Of note, the subcellular nature of spatial representation in CA3 is understudied and all the more vital given recent studies postulate weighted spatial input from CA3 may drive plasticity necessary for context dependent formation of place cells in area CA1 (*21*). With this chronic preparation we were able to track the same pyramidal cells along with their dendrites across several days (e.g., Fig 4A) and varied spatial environments. Thus, we could examine how dendrites in CA3 represent space and the nature of their input-output coupling relationship with the soma. We trained mice to run for sucrose rewards randomly distributed along the track (*27, 28*), and subsequently exposed the mice to a textured treadmill belt for a period of 12 days to elicit place cell activity. We performed two-photon imaging from days 5-12 of exposure to the same environment (belt A, Familiar, F) in 8 mice (days 1-12 in 3 mice). Somatic and dendritic segments were manually labeled, and dendrites were assigned to parent soma based on visual inspection (0-21 dendritic branches imaged per cell; see Methods for segmentation procedure). This connectivity was verified using a shuffle analysis (Fig S2-3). Calcium (Ca^2+^) activity was monitored by changes in fluorescence (df/f) (*31*) for each segment. Not all calcium events were global to all compartments of a cell (apical, basal, somatic). Many events were observed in individual dendritic compartments unaccompanied by a time-locked event in the soma. These Ca^2+^ transients must be locally generated in the dendrite and not propagated strongly enough to the soma to elicit an action potential (Fig 1E).

We then looked for spatial tuning of both dendrites and somata. Based on their calcium activity, they were classified as spatially tuned or non-tuned (*32*) using a criterion combining spatial information (*29*) and tuning specificity (*28*). We found that 41% of imaged dendrites of CA3 pyramidal neurons (43% of 1356 apical dendrites, 29% of 1210 basal dendrites, p = 0.5776, Mann Whitney test) visible in the field of view show calcium events that are stably tuned to space across laps in a session (Fig 2B-C). We call these spatially tuned dendrites “place dendrites”. We found no differences in spatial tuning quality between apical and basal dendrites, although apical dendrites are slightly more active than basal dendrites (Fig S4, median AUC, apical = 63.97, basal AUC =49.03, p < 0.0001, Mann-Whitney test). One may expect to find place dendrites in cells with spatially tuned soma, as place cells have been shown to fire in bursts (*5, 15, 33*) and such burst firing drives strong back-propagating action potentials into the dendrites (*34*).

**Figure 2:**
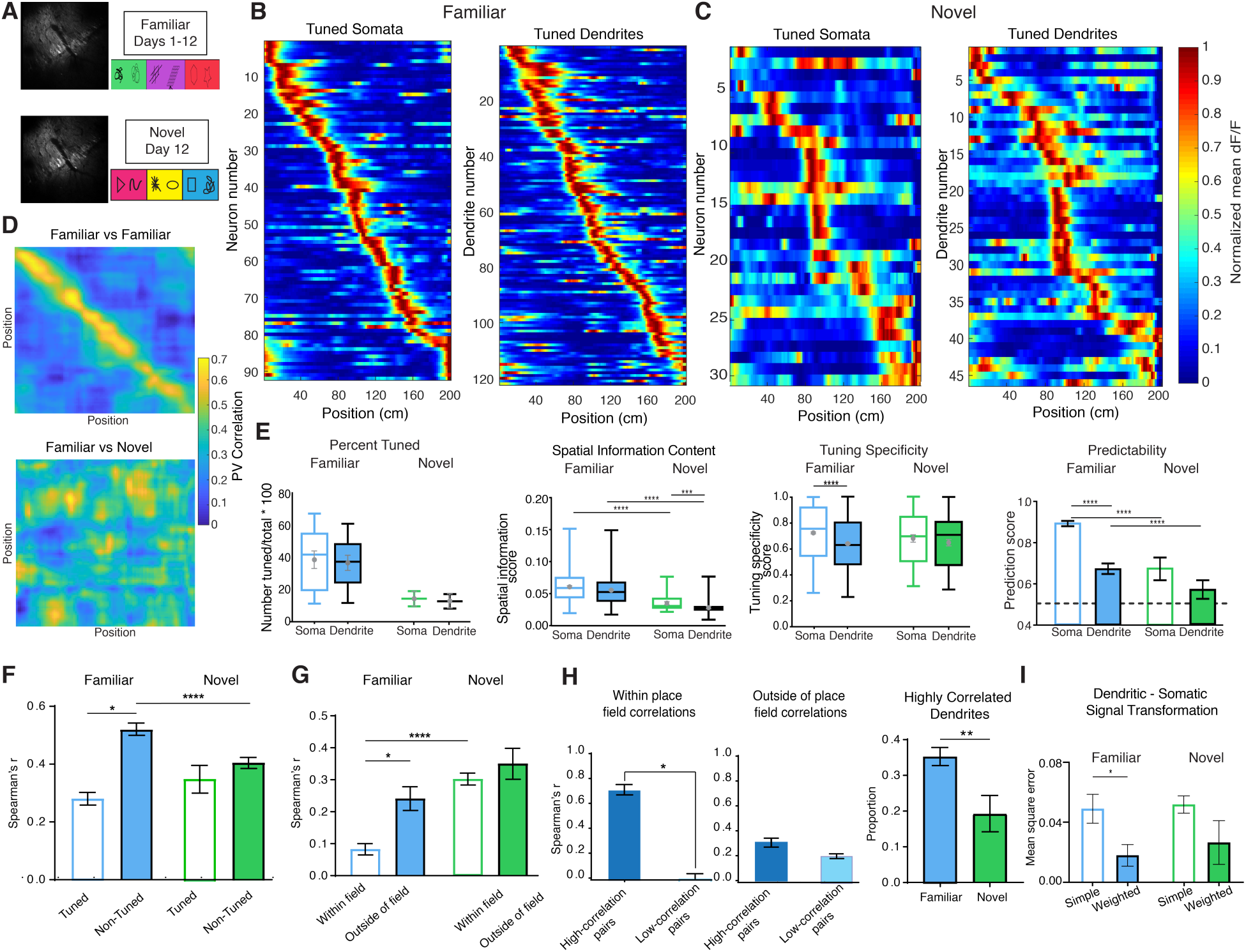
Characterization of spatial tuning: soma-dendrite decoupling in place cells in a familiar environment. **A.** Schematic showing experimental paradigm. **B.** Spatial tuning heatmaps for tuned somata and tuned dendrites in the familiar environment. Data from either compartment provides a thorough tiling of the space. Cells are ordered according to the location of their place fields. **C.** Spatial tuning heatmaps for tuned somata and tuned dendrites in the novel environment. **D.** Population vector correlation for two familiar sessions (top) and a familiar and novel session (bottom), showing a considerable degree of remapping in the novel environment. **E.** Spatial tuning metrics for somata and dendrites in familiar versus novel conditions. Far right: Ability of somata and dendrites to predict whether a soma is a spatially tuned. **F.** Spearman’s correlation scores between soma dendrite pairs for soma that are either tuned or non-tuned in familiar vs. novel environments. **G.** Spearman’s r correlation scores for tuned soma and their dendrites within the place field of that soma and outside of the place field of that soma in both familiar and novel environments. **H.** Left: Separating the within field correlation bar in the familiar environment (empty blue in G) into two groups highly correlated pairs (soma-dendrite pairs that had a Spearman’s r of 0.7 or greater throughout the session and shared a place field) and lowly correlated pairs (soma-dendrite pairs that had a Spearman’s r below 0.7 throughout the session) Middle: Separating the outside of field correlation bar in the familiar environment (filled blue in G) into the same two groups. Right: Greater proportion of highly correlated dendrites in familiar environments as compared to novel **I.** Mean square error obtained by fitting the somatic tuning curve as a weighted or a simple sum of the dendritic tuning curves for familiar and novel environments. A weighted sum of dendritic spatial tuning curves is a statistically significant better match for somatic tuning curves than a simple average in familiar environments, but not novel. Data that are not normally distributed are plotted as boxplots with a point at the mean and error bars around that point for the SEM. Normally distributed data are plotted as bar graphs. (** denotes a p < 0.01, *** denotes a p < 0.001, **** denotes a p<0.0001)

Contrarily, both apical and basal dendrites could have spatial tuning independent of their soma in that: either soma and dendrite are tuned to different spatial locations, or a dendrite is tuned to space when its soma is not (Fig 1F). Out of all place dendrites, 79% shared the same place field as their connected soma, while 21% of place dendrites were tuned to an independent spatial location. Thus, dendrites do not simply reflect the back-propagated spatially tuned activity of their soma, but can indeed hold forward-propagating, independent spatial information. This opens up the possibility that different dendrites of the same cell could hold separate spatial information. We found that branch-specific tuning does occur in CA3 pyramidal cells (Fig 1G), the first evidence for branch selectivity reported in the hippocampus. Although we separated Ca^2+^ signals from overlapping somata and dendritic segments (*31*), we wanted to verify that the branch selectivity we observed was not an outcome of improper calcium signal de-mixing and inability to discern overlap in the z-plane in our densely labeled data set. We thus validated this finding in an independent data set where we expressed GCaMP6f very sparsely in area CA3, such that individual neurons and their dendritic arbors were labeled with no overlap between cells (Fig 1H, S5). Data from this group showed similar spatial tuning metrics to the densely-expressing set (Fig S6-7). In this sparsely labeled data set where we could image and connect 104 dendritic branches to their somata (5 cells) unambiguously, we found all of the cells display branch-specific spatial tuning during at least one session. This provides *in vivo* evidence not only for locally generated dendritic Ca^2+^ events in the hippocampus but also for decoupling of dendritic activity from the somatic activity in terms of feature selectivity.

Our data showing branch-specific tuning is in direct contrast to the idea of branch spike prevalence, whereby many dendritic branches are homogenously tuned (co-tuned) to the same spatial location, that has been previously reported (*26*). These two can coexist if we consider their change with time, whereby backpropagated action potentials (bAPs) from a tuned soma may homogenize dendritic activity over time (*35, 36*). In fact, intracellular somatic recordings show that the spatial tuning of inputs became more homogenous the more an animal explored an environment (*24*). We thus hypothesized that branch-specific tuning should decrease in the place cell population with experience. To examine this, we compared the proportion of neurons that show branch-specific tuning (at least two dendrites with independent place tuning) on day 1, when the environment is novel, versus day 12 when the environment has become familiar. As we expected, the proportion of spatially tuned place cells exhibiting branch-specific tuning declines from 55% on day 1 of exposure to an environment to 19% on day 12 (p<0.05, Mann-Whitney test) of exposure to the same environment. This value is unchanged for non-tuned cells (62% on day 1, 70% on day 12, p = 0.5254, Mann-Whitney test) (Fig 1I), suggesting that subcellular organization of spatial tuning changes specifically in place cells with experience.

Branch-specific encoding of spatial information likely contributes to efficiency of spatial representation wherein *de novo* place cells exist upon first exposure to a novel environment (*11-14*). To examine how spatial tuning in dendrites might differ with novelty, we imaged the same field of view and compared tuning features of place somata and dendrites as animals randomly foraged in familiar (F) versus novel (N) track belt environments For this, we imaged the same field of view in a novel environment (belt B) one hour after imaging in the familiar environment on day 12. Whether we used somatic or dendritic activity, we observed place ensembles that spanned the entire length of both the familiar and novel treadmill tracks (n = 631 tuned somata, 907 tuned dendrites) (Fig 2B-C). This shows that as an ensemble, dendritic branches can independently represent the entire spatial environment. We compared the population vector (PV) correlation of somatic activity on two different days of exposure to the familiar environment (0.4302) to the PV correlation between activity in the familiar vs. novel environments on day 12 (0.0415) (Fig 2D) and found a significant drop in PV correlation (p <0.0001, Mann Whitney test). These results validate that the spatially tuned activity of the CA3 place cell population is sufficiently stable across days within the same environment, while introducing the animals to a novel spatial environment triggers an extensive reorganization of the place cell ensemble. This is in agreement with the vast literature showing distinct neural representations of different environments (*37*)

While there exists a distinct representation of the novel spatial environment from the early trials of the session, we found that the proportion of spatially tuned dendrites and soma are both reduced in the novel compared to familiar environment (*somata*, F: 41%, N: 14%; *dendrites* F: 37%, N: 13%) (Fig 2E). Next, we compared activity and spatial tuning properties of dendrites versus somata in these two conditions while controlling for number of laps run and speed of the animal (Fig S9A). Both place somata and place dendrites have higher activity rate in the familiar environment compared to the novel environment (mean AUC/min: *somata*, F: 126.7, N: 86.04, p<0.001; *dendrites*, F: 87.5, N: 46.83, p<0.0001, Mann-Whitney test with Bonferroni correction for multiple comparisons) (Fig S8). In terms of spatial tuning metrics (Fig 2E), both soma and dendrites showed drastically reduced spatial information content in the novel environment compared to the familiar (*somata*, F: 0.059, N: 0.031 p < 0.0001; *dendrites*, F: 0.053, N: 0.027, p < 0.0001; Kruskal-Wallis test with multiple comparisons) but no change in their tuning specificity (median: *somata*, F: 0.62, N: 0.70, p = 0.1288;*dendrites*, F: 0.75, N: 0.69, p = 0.7391; Kruskal-Wallis test with multiple comparisons) or place field width (median *somata* F: 17.68 cm, N: 18.68 cm, p = 0.0649; *dendrites*, F: 18.45 cm, N: 19.08 cm, p = 0.3334, Kruskal-Wallis test with multiple comparisons) (Fig S9).

Beyond the context-related differences, our data provide the first glimpse of how dendrites themselves render a representation of the spatial environment. Thus, we closely examined how the activity and place tuning properties of the CA3 dendritic place ensemble compares to that of the somatic place ensemble within each condition (Familiar and Novel). We observed that somatic activity was significantly higher than their dendritic counterparts in both familiar (p<0.0001) and novel (p<0.01) environments (Fig S8). We found that while in both conditions dendritic place fields are as precise as those exhibited by somata in their place field width (dendrites vs. somata, F: p = 0.05, N: p = 0.8083) (Fig S9), place dendrites have slightly poorer spatial information score (p < 0.001, dendrites vs. somata) in the novel environment and tuning specificity (p<0.0001) in the familiar environment.

Previous studies have implicated that dendritic activity is predictive of feature selectivity in both cortex (*16*) and hippocampus (*22, 35*). To directly test the accuracy with which the calcium activity of dendrites could predict the spatial tuning of their corresponding soma, we used our data to train a machine learning algorithm. We trained an Artificial Neural Network (ANN) classifier using the mean calcium event profiles across space (of dendrites or somata) as input to predict the label (place or non-place) of each cell (Fig S10). We then tested the classifier on an unseeded dataset and compared its prediction accuracy to the true labels (see Methods). We found that dendritic activity could indeed predict somatic tuning in the familiar condition (dendrites AUC: 0.68±0.019), albeit with significantly lower accuracy than their soma (soma AUC: 0.86±0.013, p<0.0001; two-way ANOVA with multiple comparisons, Bonferroni’s correction). While somatic activity remained predictive of spatial tuning in the novel environment, albeit with low accuracy, dendritic activity could no longer predict somatic spatial tuning (Fig 2E) (soma: 0.69±0.063 AUC, dendrites: 0.50±0.094 AUC; p<0.0001; two-way ANOVA with multiple comparisons, Bonferroni’s correction). Together, these data show lower quality spatial tuning in a novel environment as compared to a familiar. This suggests a possible role for experience-dependent sculpting of inputs to improve the quality of spatial tuning over time. Repeated exposure to the same environment can foster coincident activity dependent long-term plasticity that results in strengthening of spatial inputs (*24*). In contrast, one-time exposure to a novel environment may be insufficient to support mechanisms that boost spatial inputs and filter out non-spatial “noise”.

In area CA1, *in vivo* current injection of a dendritic spike like waveform into the soma of silent cells or even place cells at any location on the track belt triggers the formation of a place field at that given location (*21*). This supports our finding that a cell may encode multiple spatial locations through dendritic branch-specific spatial tuning. It also implies that spatial tuning selectivity of a soma likely arises as a function of selective propagation or coupling of dendritic input to the soma rather than specificity of overall input content to the cell. Even in CA3 place cells that only have one place field, we show evidence for dendritic branch-specific tuning (Fig 1G-H). If all selectively tuned dendritic branches propagated to the soma, that soma would appear untuned - firing at many different locations in space. Thus, we hypothesized that for the development of a high-fidelity place cell, signal flow from dendrites to soma has to be selectively sculpted.

To examine this, we first investigated the relationship between dendritic and somatic activity of place cells versus non-place cells. From the same dataset as above, we computed correlation scores (Spearman’s r) between calcium traces for each dendrite and its corresponding soma across the entire session. We found a very low correlation between dendritic and somatic activity in tuned cells (median r = 0.3536 for tuned, n = 450 tuned cells) as compared to non-tuned cells in the familiar environment (median r = 0.5910 for non-tuned, n = 230 p<0.0001, Mann-Whitney test, two-tailed) (Fig 2F, left). These correlation relationships might arise from higher activity in non-tuned cells than tuned cells, simply increasing the statistical probability of correlation. In fact, we found the exact opposite in our data: tuned cells show greater activity in both their somata and dendrites than non-tuned cells, indicating that activity rate is not a confound. (Fig S8, somatic mean AUC/min: *tuned:* 72.76, *non-tuned cells:* 41.73, p < 0.0001; dendritic mean AUC/min *tuned cells:* 52.97, *non-tuned cells:* 45.90, p = 0.0161, one-way ANOVA). To assess whether this low correlation in tuned cells was an artifact of comparing dendritic activity from outside the somatic place field, we extracted correlation scores for time windows only when the animal was within the place field of the soma.

Surprisingly, this decoupling was further exacerbated within the place field of the soma, a time window when we expected the soma to be most receptive to all dendritic activity to allow for strong spatial tuning (*26*) (median r = 0.1575 for within field, r = 0.3672 for outside of field, p<0.0001, Mann Whitney test, two-tailed) (Fig 2G, left). On closer examination this low correlation is quite intuitive because we are examining the correlation from all dendrites, even those that are either untuned or independently tuned place dendrites. Thus, we next looked at only soma-dendrite pairs that showed high correlation throughout the session (Spearman’s r of at least 0.7, n = 171). In all cases these were dendrites that were tuned to the same spatial location as their soma. The in-field correlation for these pairs was very high (median r = 0.7985) while the in-field correlation for all other dendrite soma pairs (n = 279) was nearly zero (r = 0.0227) (Fig 2H, left), suggesting that there is indeed open signal flow between the soma and select dendrites active only during a specific spatio-temporal window (p<0.0001, Mann Whitney test, two-tailed). We wondered if this decoupling from most dendrites was a feature that predisposed cells to become place cells, or was something that developed over time as cells became tuned. To answer this question, we looked to the novel environment, where new place cells are formed rapidly. Interestingly, the difference in dendrite-soma coupling between tuned and non-tuned cells observed in familiar environments was not present in the novel environment (median r = 0.3243 for tuned, 0.4155 for non-tuned, n = 54 for tuned, n = 406 for non-tuned p = 0.4951 for novel) (Fig 2F-G, right). Moreover, there was a significantly higher proportion of dendrites that showed high correlation with their soma in the familiar environment (0.3538) than in the novel (0.1935, p = 0.0099) (Fig 2H, right). This suggests that over time, as an animal familiarizes itself with an environment, a place soma is isolated from signal flow from the majority of its dendrites, while developing a strong correlation in its activity with select place dendrites.

To validate the idea that inputs from different dendrites propagate with different efficiencies, we next compared whether weighted averages of dendritic activity were closer to somatic activity of a given cell than simple averages. We performed a linear regression on the dendritic tuning curves to find the optimal weights to minimize the error between the weighted sum of the dendritic traces and the corresponding somatic tuning curves for each cell. We performed this analysis only in our sparsely-expressing dataset where we could unambiguously assign a large number of dendrites to their corresponding soma. In familiar environments, the resulting errors in the weighted average set were significantly lower than those derived from a simple averaging of the dendritic curves (weighted = 0.0174, simple = 0.0484, p = 0.0460, Wilcoxon test) (Fig 2I, left) consistent with heterogeneous propagation efficiencies from dendrites to soma. In novel environments, the weighted average did not significantly out-perform the simple average (weighted = 0.0267, simple = 0. 0520, p = 0.25, Wilcoxon test) (Fig 2I, right). Although more data might reveal a significant difference in novel environments, our results indicate a more homogeneous propagation than in familiar environments.

Next, we simulated multi-compartment CA3 neurons with dendritic tuning properties constrained by our data (Fig 3A, see methods). To simulate familiar environments, well-tuned dendritic activity propagated to the soma with a non-uniform propagation efficacy such that some dendrites were more strongly connected with the soma than others. In novel environments, dendrites were only weakly tuned, and propagation efficacy was more homogenous. Under both conditions, dendritic and somatic place fields spanned the entire environment (Fig 3B-C), with noisier place fields in novel environments. The emergence of somatic place fields from diversely tuned dendritic place fields suggests two contributing mechanisms: a bias in dendritic tuning to a specific location and non-uniform dendrite-to-soma propagation efficiency (Fig S11).

**Figure 3:**
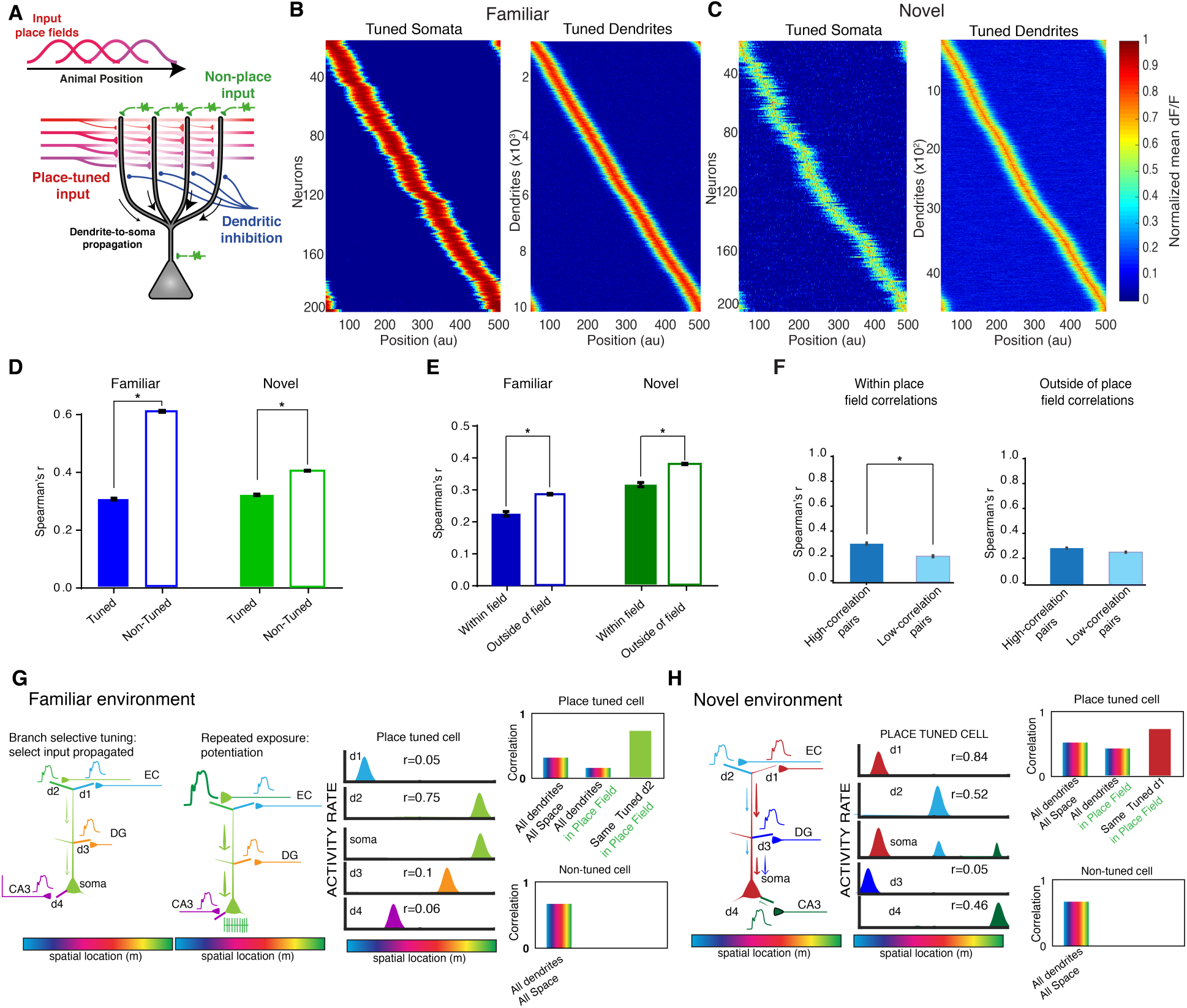
Multi-compartment modeling of spatial coding in CA3 place cells. **A.** Cartoon of CA3 multi-compartment model. **B.** Spatial tuning heatmaps for the model generated data in the familiar environment. **C.** Spatial tuning heatmaps for the model generated data in the novel environment. Analogous analysis to A, using modeling generated data. **D.** Analogous analysis to Fig 2E, using modeling generated data. **E.** Analogous analysis to Fig 2F, using modeling generated data. **F.** Analogous analysis to Fig 2G, using modeling generated data. **G.** Schematic of potential mechanism of correlation relationships observed in the familiar environment. A proposed role for gating forward and potentiation of “preferred” input in a familiar environment. **H.** Schematic of potential mechanism of correlation relationships observed in the novel environment whereby potentiation has not yet developed. (*** denotes a p < 0.001, **** denotes a p<0.0001)

**Figure 4:**
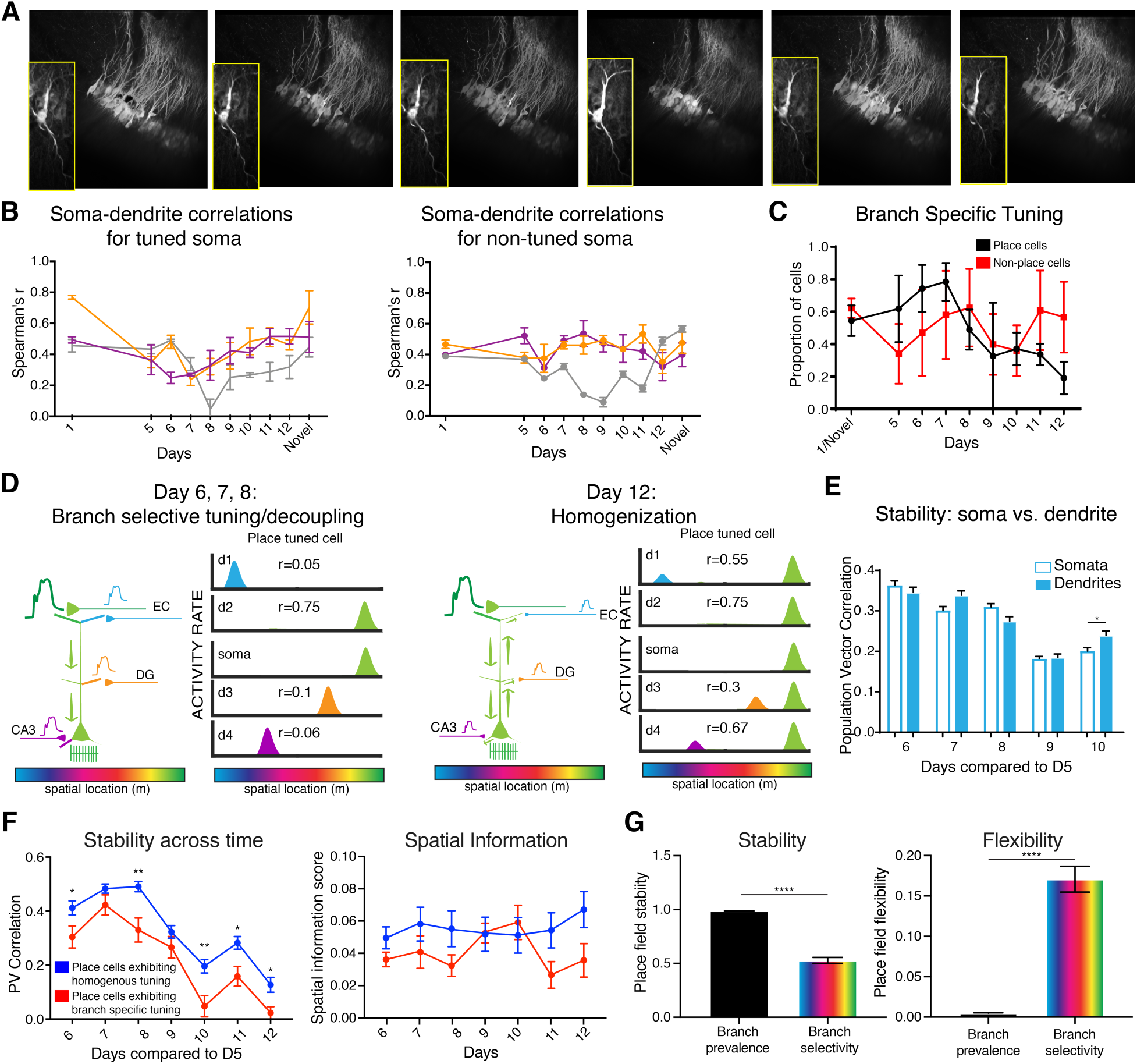
Experience-dependent changes in tuning. **A.** The same field of view and an example cell across 6 days (days 5-10 for this example). **B.** Tracking the correlation between soma and dendrites across time for tuned soma (left) and non-tuned soma (right). There is an observed trough in the data from the tuned soma, with the day of lowest correlation varying by animal. **C.** Proportion of cells that show branch-specific tuning across time. Of note, this curve shows an inverse relationship to the soma-dendrite correlations plotted across time. **D.** Schematic of potential mechanism of correlation relationships observed across time. With increased familiarity, backpropagating activity homogenizes tuning of cell compartments. **E.** PV correlations across time with day 5 as a reference show no differences between stability of place soma and place dendrites. **F.** Place cells that exhibit homogenous tuning on day 5 are more stable across time than cells that exhibit branch-specific tuning, showing that branch selective tuning may confer a propensity for plasticity. Spatial information is unchanged across time, indicating that the decrease in stability is not a byproduct of worse spatial tuning. **G.** Place field stability and (left) and flexibility (right) for simulations under branch prevalence and branch selective tuning conditions. Branch selective tuning confers flexibility to the spatial code. (* denotes a p <0.05, ** denotes a p < 0.01, **** denotes a p <0.0001)

We then used our simulations to investigate the mechanisms yielding the observed dendrite-soma correlations. By controlling the level of non-place tuned inputs, our model reproduced the levels of correlation in both novel and familiar environments (Fig 3D-F, see also Fig S11 and Methods). When compared to familiar environments, the difference between tuned and non-tuned cells was smaller in novel environments, but significant (Fig 3D, right). The difference between simulation and experiment in novel environments might be a reflection of the higher number of samples in our simulations compared to the number of samples in our experiments. Interestingly, the higher correlations outside of place fields were a direct result of a global, non-spatial input delivered equally to all compartments in our neuron model. If these “non-place” inputs were removed, soma-dendrite correlations became higher within place fields than outside of place fields (Fig S11). These combined experimental and modeling results suggest an experience-dependent change in the coupling between place soma and their dendrites. Based on these results, we posit that from many place dendrites, only activity of a few “preferred” dendrites is propagated forward, and with repeated somatic coactivity, this input is perhaps potentiated via plasticity mechanisms over time. This could lead to the observed decoupling between most “non-preferred” dendrites of place-tuned soma that is seen initially (Fig 3G). In the context of many tuned dendrites, blockade of signal flow from “non-preferred” spatial inputs could be achieved via depression as bAPs from burst-firing tuned somata repeatedly occur out of sync with the forward propagating, differentially tuned signal in the dendrite; or from branch-selective inhibitory gating of “non-preferred inputs” (*38*). On the contrary, in a novel environment, perhaps plasticity has yet to occur and thus the decoupling is absent (Fig 3H).

To further examine this experience-dependent reorganization, we tracked the same dendrite-soma pairs across all 12 days (Fig 4A) and computed correlations between each pair on each day. We found that on day 1 of exposure to an environment (when it is novel), tuned soma showed similar correlation with their dendrites as was seen in the novel data above (0.4823 on day 1, 0.5003 in novel, p = 0.8158, Mann-Whitney test). Upon repeated exposure to the environment across several days, this correlation dropped for all animals until it reaches a trough on day 5, 6, or 7. After this trough was reached, the correlation began to rise again. In a novel environment that was experienced on day 12, the correlation was again high relative to the values seen in the familiar environment (Fig 4B). Interestingly, the prevalence of branch-specific tuning across time bore an inverse relationship to these correlations (Fig 4C). An increased prevalence of branch-specific tuning coincided with a decreased correlation between soma-dendrite pairs and vice versa. This is consistent with our conceptual framework in which dendrites tuned to a different location than their somata need to be decoupled from the soma to maintain fidelity of spatial tuning output of that cell. (Fig 4D, left). Over time, however, the depression of non-specific inputs would render a cell more homogenously tuned (Fig 4C-D, right) and the average soma-dendrite correlation would be increased, as is seen on days 7-12 in our data (Fig 4B, left). Another possibility that could lead to the changes we observe in correlation and branch-specific tuning across time is a decrease in stability of dendritic activity. To rule this out, we computed PV correlations of the activity of tuned somata and dendrites across time comparing the activity for each day to that of day 5 (Fig 4E, S11). We found that place somata and place dendrites show no differences in stability, except for a single day in which the dendritic ensemble showed slightly greater stability than the somatic ensemble (PV corr *dendritic:* 0.2395, *somatic:* 0.2008, p < 0.05, Kruskal-Wallis test with multiple comparisons). This indicates that a decline in dendritic stability is unlikely to be the cause of the observed decrease in branch specific tuning across time.

What is the benefit of branch-specific tuning if it just dissipates over time? Is it simply an intermediate step for achieving spatial tuning of a cell or does it play a specific role in shaping ensemble dynamics? Some place cells remain highly stable across time, while others are more flexible. Previous studies show that increased homogeneity of dendritic activity leads to more stable, precise place fields (*26*). Conversely, does the degree of flexibility depend on the ability of a cell to represent multiple spatial locations within its dendritic arbor? To examine this, we split the somatic data by whether the cell exhibited branch-specific tuning or homogenous tuning. We found that the homogenously tuned cells showed greater stability across time (Fig 4F, left) as compared to cells exhibiting branch-specific tuning. To test if the greater drop in PV correlation in the branch specific tuning set was due to a degradation in the quality of tuning across time, we quantified their spatial information content across days 6-12 and found no accompanying change (Fig 4F, right). Taken together this means that although cells may change their place fields from day-to-day (contributing to the drop in PV correlation), they are equally well tuned across familiar days (as seen by consistently high spatial information score). This suggests that branch-specific tuning may confer flexibility to the map of cognitive space without degrading the quality of the cognitive spatial map. We explored this idea by simulating two groups of cells, one in which cells exhibited branch-specific tuning, and one in which dendrites were homogenously tuned (deemed “branch prevalence” dominated). We then compared these two conditions and assessed their effect on place field stability and flexibility. As we expected, the model neurons with homogenously tuned dendrites showed greater place field stability, but were inflexible to change (stability = 0.98, flexibility = 0.005) (Fig 4G, left). The model incorporating branch-specific tuning was flexible enough to remap (stability = 0.53, flexibility = 0.17) (Fig 4G, right), a phenomenon observed widely in CA3 (*14, 30, 37, 39*). Branch specific tuning could thus provide a dynamic range of spatial feature selectivity to place cells enabling rapid remapping (as seen by flexible place tuning) for otherwise static representations of space.

In summary, we find high fidelity place dendrites in area CA3 belonging to both tuned and non-tuned somata. When place dendrites belong to a tuned soma, they may encode place field locations different from their somatic place fields (although they are often tuned to the same location). We provide the first experimental evidence for hippocampal place cells exhibiting dendritic branch-specific tuning. Even in neurons exhibiting branch-specific tuning, the soma typically expresses only a single place field and thus becomes decoupled from many of its dendrites – a mechanism by which cells can maintain their specific spatial activity in the face of differentially tuned dendrites. This decoupling only occurs with repeated exposure to an environment, and as such is likely the result of experience-dependent plasticity. While we do not know the mechanism by which certain dendritic inputs are selectively gated while others are decoupled, we can speculate that coincident activity-dependent plasticity or selective inhibition may be at play. This multi-stage, compartmentalized organization of spatial information within a single hippocampal place cell can greatly increase its computational power, and thus allows for flexibility of spatial representation.

## Methods

### Cranial Window Surgery

All experiments were approved by New York University’s Animal Care and Use Committee. 4 female and 4 male mice were anesthetized using isoflurane (1.5-2.5%) and a small (0.5 mm) hole was drilled in the skull above dorsal area CA3 of hippocampus (1.6 mm lateral, and 1.4 mm caudal of Bregma). 2 females and 2 males had injections and implants on the left side, and the remaining 4 animals had the procedure on the right side to control for laterality. Injections of AAV1.CamKII.GCaMP6f.WPRE.SV40 (commercially generated at Penn Vector Core, titer: 2.76×10^13^ GC/ml, 23 nL per site) were made at two sites, three depth levels each (1.5 mm lateral, 1.3 mm caudal of Bregma; 1.7 mm lateral, 1.5 mm caudal of Bregma; depths of 1.8, 2, and 2.2 mm below the dural surface). For sparse expression, injections were made at only the 1.8 depth level using a mixture of AAV1.CamKII-Cre (commercially generated at Penn Vector Core, titer: 2.71× 10^13^GC/ml diluted 100,000 fold with aCSF) and AAV1.Syn.Flex.GCaMP6f.WPRE.SV40 (commercially generated at Penn Vector Core, titer: 1.64×10^13^ GC/ml) as in (*25*) at 23 nL per site. After injection, a 3 mm craniotomy was made with the injection site at the center. The skull was removed, and a vacuum system was used to gently remove the overlying cortex and external capsule. aCSF (NaCl 119mM, KCl 2.5 mM, NaH_2_PO_4_ 1.25 mM, NaHCO_3_ 24 mM, glucose 12.5 mM, CaCl_2_ 2mM, MgSO_2_ 2mM) at 0° was used to irrigate the area throughout the duration of the procedure. A cranial window (3 mm diameter, 1.7 mm length stainless steel cannula attached to 3 mm diameter glass coverslip) was then implanted over the area. The window was sealed to the skull using Vetbond, and a custom designed 3D-printed plastic headpost (*40)* was cemented over the skull.

### Behavioral training

Two days after cranial window implantation, water scheduling was begun. Mice were provided with 1 mL of 5% sucrose per 24 hours. A custom-built behavior system (as described below) fitted under the two-photon microscope was used to train and run mice head-fixed on a treadmill. Mice were then trained to run on a 200 cm untextured treadmill for sucrose rewards presented randomly throughout the track (one 15-20 min session per mouse per day). If mice did not receive a full 1 mL of sucrose during training, they were given the balance in water after training. Once mice reached a running speed of 2 laps/min, place cell imaging was begun. Place cell imaging was performed using a textured track with 3 macrotextures and 6 microtextures (2 microtextures per macrotexture). Treadmill belts were created using 2 cm wide ribbon (3 types to create the 3 macrotextures), and microtextures were created using Velcro pieces, foil dots, rhinestones, glue dots, and textured fabric. Each mouse ran on the treadmill for ∼15 minutes per session, 2 sessions per day for 11 days. On day 12, mice were exposed to a novel treadmill belt. To control for treadmill textures, the 8 mice were run in 3 separate cohorts, with each cohort having different treadmill belts for both familiar and novel conditions.

#### Behavior apparatus for reward delivery and behavior tracking

Briefly, the same behavior setup (as described in (*29*)) was used for training and running mice during imaging experiments. The system was equipped with i) a 3D printed head-post mounting and fixation assembly, ii) treadmill track belt with RFID tags communicating with an RFID reader (ID-20LA, SparkFun Electronics) moving on 3D printed wheels coupled to a rotary encoder (S5-720, US Digital) for position sensing, iii) a solenoid valve (Parker) driven reward delivery system coupled to a capacitive lick sensor (SparkFun Capacitive touch Breakout AT42QT1010) for detecting the licking of the mouse during collection of the sucrose water rewards. A microcontroller (Arduino Mega, Adafruit) connected to a shield (OM2, OpenMaze) was used to control the input-output for the behavior.

### Two-Photon Imaging

We used the same imaging system as described previously (*29*). Briefly in vivo two photon imaging was performed using a dual galvanometric and resonant laser scanning two-photon microscope (Ultima, Bruker), coupled to a tunable Ti:Sapphire laser (MaiTai eHP DeepSee, Spectraphysics) pulsed at a 80 MHz repetition rates and <70fs pulse width along with dispersion compensation. GCaMP fluorophore was excited at 920 nm, using a resonant scanning X-galvanometer (8kHz) paired with a 6mm standard scanning Y-galvanometer. The scanning system was mounted on movable objective Ultima microscope, equipped with an orbital nosepiece coupled to a 16X, 0.8 NA, 3 mm water immersion objective (Nikon) and a piezo drive for angled imaging and ultrafast volumetric scanning. Imaging was performed at a scan speed of 29 fps, using 512×512 frame size (1.012 µm/ pixel resolution). Fluorescence signal was separated into green channel and red channel using a dichroic mirror (T565lpxr, Chroma Technology) and emission filters (green, ET510/80m-2p; red, ET605/70m-2p, Chroma Technology), respectively. Each emission signal was detected using high-sensitivity GaAsP photomultiplier tubes (model 7422PA-40 PMTs, Hamamatsu).

### Data Analysis

Analysis was performed using the CaImAn Matlab package (*30*), ImageJ, and custom written Matlab scripts. CaImAn was used to motion correct each time-series. Both somatic and dendritic regions of interest (ROIs) were hand drawn using ImageJ. Only dendrites that could be traced to their cell body were included in the analysis. ROIs for dendrites that could not be traced to a cell body were still drawn so that their calcium signals could be de-mixed from adjacent/overlapping neuropil and somata. Calcium signals from those dendrites were subsequently excluded. Dendritic branches were defined as separate segments anytime a branch point was observed.

#### Ca2+ transient detection

Significant transients were identified as events that started (onset) when fluorescence deviated 2σ (standard deviation) from the baseline trace and ended when it returned to within 0.5σ (offset). We then recalculated the baseline of the full calcium trace after masking frames occurring during previously identified significant transients. The standard deviation was then recalculated and transients re-estimated. Transients less than 1s were removed. This procedure was repeated three times. For the spatial information and the spatial tuning calculation we restricted our analysis to running epochs, defined as consecutive frames of forward locomotion of at least 2s duration and with a minimum peak speed of 5cm/s. Consecutive epochs separated by <0.5s were merged. ROIs that were active during less than 20% of laps run were excluded.

#### Spatial information

The spatial information was computed from the smoothed rate map: a Gaussian filter (sigma = 3 bins) was applied to the calcium event count, and occupancy across bin positions and the smoothed rate map was constructed by dividing the smoothed event count by the smoothed occupancy. For each cell the spatial information content (*41*) was calculated according to the formula

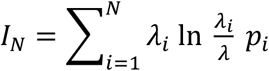

Where, *λ*_*i*_ and *p*_*i*_ are the transient rate and probability of occupancy of bin *i*, λ is the overall transient rate, and *N* is the number of bins. We computed *I*_*N*_ for multiple values of N = 2, 4, 5, 8, 10, 20, 25, and 100. We then created 100,000 random reassignments of the transient onset times within the running-related epochs and re-computed the values of 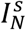, where *s* is the index of the shuffle. This measure is biased by both the number of bins chosen and by the number of events fired by the cell. Performing shuffles on a per-cell basis addresses the latter bias. To roughly correct from the bias associated with binning, we subtracted the mean of this null distribution from all estimates to obtain values 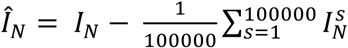. Finally, we computed a single estimate of the information content for the true transient onset times, 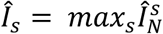 and for the shuffles,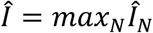. The spatial tuning p-value was taken as the fraction of values of *s* for which *Î* exceeded *Î*_*s*_. A cell was considered to have significant spatial information content if p<0.05 (*28*).

#### Tuning specificity

Spatial tuning vector was calculated as 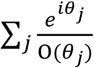

Where *θ*_*j*_ is the position of the mouse at the onset time of the Ca2+ transient *j*, O(*θ*_*j*_) is the fraction of frames acquired at position *θ*_*j*_. We assessed the significance of the spatial selectivity for each cell by generating a null distribution of shuffled transient onset times and recalculating the tuning specificity 10,000x. The p value was defined as the fraction of this null distribution that exceeded the neuron tuning specificity.

ROIs, regardless of whether they were somata or dendrites, were classified as spatially tuned if they met both spatial information and tuning specificity criteria.

#### State-dependent Correlation (Fig S2 Panel A)

This analysis was done on a subset of the data from 4 mice. Continuous ΔF/F traces for soma and dendrite were split into periods of mobility (treadmill speed > 2 cm/s) and immobility (speed <= 2 cm/s), and the Pearson’s correlation coefficient was calculated between traces of soma-dendrite pairs in each condition.

#### Matched vs Unmatched Correlation (Fig S2 Panel B)

The correlation between matched soma-dendrite pairs was computed as described above. To generate unmatched correlations, the correlation coefficients between a soma and all dendritic ROIs that were not identified as belonging to that soma were computed. This was done for all soma in a session, with the results aggregated together. All correlations were computed only using data during mobility.

The distance between a soma-dendrite pair (matched or unmatched) was the Euclidean distance between the centers of each ROI.

The moving average for matched and unmatched pairs was computed as the mean of data points falling into 50 bins with edges logarithmically spaced between 4 and 560 cm. This moving average was then smoothed with a Gaussian kernel with σ = 1 bin. Only bins with at least 2 data points in a given population were plotted.

#### Coactivity Analysis (Fig S2 Panel C)

All analysis was done only using data during mobility. Each ΔF/F trace was z-scored with respect to itself and identified as “active” if the z-scored value exceeded 2. For each soma-dendrite pair, a coactivity score was computed as the following ratio:

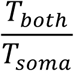

Where T_*both*_ is the amount of time both the soma and dendrite were simultaneously active, and T_*soma*_ is the amount of time the soma was active, regardless of the activity state of the dendrite. This yields a bounded measure between 0 and 1, with 0 indicating a complete absence of simultaneous activity in the two ROIs, and 1 indicating total synchrony.

We used a bootstrapping procedure to estimate the null coactivity between a soma-dendrite pair, defined the amount of coactivity between the pair expected by chance. The ΔF/F trace of the dendrite was first time-reversed and then circularly shifted by a random amount (at least 1 second) to obtain a null dendritic trace. The coactivity between the soma and this null dendritic signal was then computed. This was repeated 100 times with different shift amounts, and the null coactivity was defined as the 95^th^ percentile of the resulting distribution. Thus, if the coactivity of a soma-dendrite pair exceeded the null coactivity so defined, then the pair exhibited statistically significant coactivity at the p<0.05 level.

#### Branch specific tuning classification

Only cells that had two or more dendrites visible in the field of view were considered for branch-specific tuning analyses. A cell was considered to display branch specific tuning only if one or more dendrites displayed in the field of view were tuned to different locations than the remaining dendrites.

#### Machine learning classification

We used classification methods to classify pyramidal neurons into place cells vs. non-place cells, based on the calcium activity of either the soma or selected dendrites. We assessed different classifiers (e.g., k-Nearest Neighbors, Naïve Bayes, Random Forests, Quadratic Discriminant Analysis, Logistic Regression, Artificial Neural Networks (ANN), etc.) and chose the one achieving highest performance under control conditions, when using the somatic recordings as input (the ANN). To evaluate the models we split the dataset into training and test sets (80-20) and performed an exhaustive grid search for various hyperparameters (Table S1) using 5-fold Cross Validation. Classification was performed using the MLPClassifier() function in the Scikit-learn(*42*) module in python. MLPClassifier is a traditional Artificial Neural Network classifier (*43*). The network consists of two hidden layers with 200 and 100 nodes, respectively. The algorithm used for learning is the stochastic gradient descent with batch size equal to 100. We used adaptive learning rate, with initial value 0.2 and decreased by 5 if the loss remained constant for two consecutive epochs. These parameters were kept constant across all runs. We analyzed two experimental conditions: (i) familiar, and (ii) novel environment. The data were split in training and testing data sets under the constraint of equally represented classes. To achieve class equality, a randomly chosen portion of the data was left out. Input was provided in a matrix form, where columns represent neurons and the rows represent the corresponding mean calcium event rate (see experimental methods for a description of event rate). Input matrices were different for somatic and dendritic calcium signals. For dendritic signals, in cases where more than one dendrite was provided, we used the activity of the dendrite that had the highest correlation with the soma (Pearson correlation coefficient). Labeling (place cells vs. non-place cells) was provided as a vector with one and zero, values, respectively. Zero-padding was used to create same-length vectors. The classifier was executed 1,000 times, with randomly partitioned training/test sets. The above training/testing process was done for each condition (control / novel) and each type of data (somatic vs. dendritic), separately. Performance was assessed using the area under the receiver operating characteristic (ROC) curve (AUC)(*44*). Plots report the mean AUC value across repetition trials and error bars indicate standard deviations. To compare the performance of the classifier amongst different groups (i.e., familiar vs. novel environment and soma vs. dendrite), AUC scores were analyzed using two-way analysis of variance (ANOVA) followed by post-hoc tests using the Bonferroni method whenever statistical significance was observed. The α was chosen to be 0.001 for statistical significance. To correct for multiple comparisons, the α was divided by the number of tests according to the Bonferroni procedure

**Table S1.** Hyperparameters for classifiers tested.

| Classifier | Hyperparameters |
| --- | --- |
| Logistic Regression (LR) | Regularization strength ('C') |
| Linear Discriminant Analysis (LDA) | • Solver method ('solver') |
| Quadratic Discriminant Analysis (QDA) | • Covariance regularization ('reg_param')<br>• threshold for rank estimation ('tol') |
| K Nearest Neighbors (KNN) | • Number of neighbors ('n_neighbors')<br>• Distance metric ('metric') |
| Decision Trees (DT) | • Function to measure the quality of a split ('splitter')<br>• Strategy used to choose the split at each node ('criterion')<br>• Number of features when looking for the best split ('max_features') |
| Gaussian Naive Bayes (GNB) | • Portion of the largest variance of all the features that is added to variances for stability ('var_smoothing') |
| Support Vector Machine (SVM) | • Regularization parameter ('C')<br>• Coefficient for 'rbf' kernel ('gamma') |
| Random Forest (RF) | • Number of trees in the forest ('n_estimators')<br>• function to measure the quality of a split ('criterion')<br>• Whether bootstrap samples are used when building trees ('bootstrap') |
| Artificial Neural Network (ANN) | • Number of layers and nodes per layer ('hidden_layer_sizes')<br>• Activation function in the hidden layer(s) ('activation')<br>• Initial learning rate ('learning_rate_init') |
| AdaBoost (ADB) | • Maximum number of estimators at which boosting is terminated ('n_estimators')<br>• Shrinkage of the contribution of each classifier ('learning_rate')<br>• Algorithm to learn the parameters ('algorithm') |

#### Dendritic-somatic signal transformation

Using our sparsely-expressing dataset, we tested whether the somatic activity could be approximated by a simple average of the dendritic activity and whether an weighted average would improve this approximation. The approximated somatic activity was given by

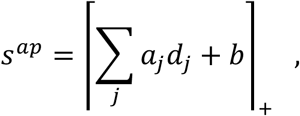

where *d*_*j*_ is the activity of dendrite *j, a*_*j*_ is a positive coefficient, b is a number, and ┌·┐_P_ is a rectification function that sets all negative values to zero. The values for the coefficients *a*_*j*_’s and *b* were chosen to minimize the mean square error between the approximated somatic activity and the activity observed experimentally. For a simple average, the coefficients were constrained such that *a*_*j*_ = *a* for all dendrites.

### Modeling

#### Neuron model

We simulated CA3 pyramidal cells using a multicompartment, rate-based model. The neuron model was composed of 100 dendritic compartments that projected to one somatic compartment. Each dendritic compartment *i* was modelled as a rate-based unit and described by a “potential” variable 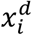 whose temporal evolution follows

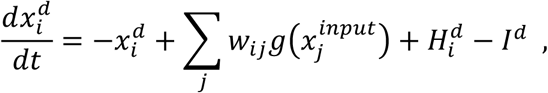

where 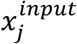 is the potential variable for input neuron *j, w*_*ij*_ is the weight of the synaptic connection from input neuron *j* to dendritic compartment *i*, 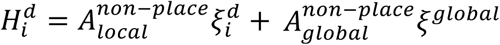 is a noise term (non-place inputs), *I*^*d*^ is the dendritic inhibition, and *g*(*x*) is the unit’s instantaneous activity, measured relative to a low level of spontaneous activity. The function *g* is given by

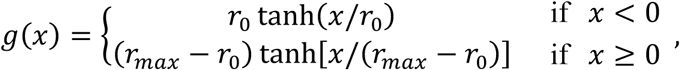

with baseline activity *r*_0_ = 1, and maximum activity *r*_*max*_ = 20. Therefore, the activity of the dendritic compartment *i*, relative to the baseline activity, is 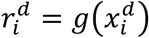.

The somatic potential for each simulated neuron was given by a similar equation,

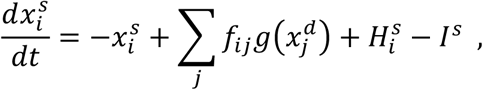

where *f*_*ij*_ is the efficiency of the dendrite-to-soma propagation from dendritic compartment *j* to soma for neuron *i*, 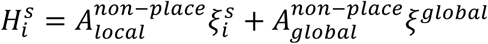 is a noise term, and *I*^*s*^ is the somatic inhibition. The somatic activity for neuron *i*, relative to baseline, is 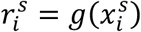.

The noise terms, 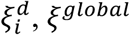 and *ξ*^*s*^, were modelled as independent Ornstein-Uhlenbeck processes. The local terms, 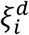 and *ξ*^*s*^, with time constant 100 ms, and the global term, *ξ*^*global*^, with time constant 10 ms. The terms 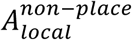 and 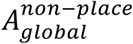 are the amplitudes of the independent (local) and global non-place inputs to all the compartments. The values for 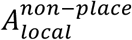 and 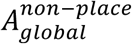 were chosen to fit our experimental data such that the mean within field, outside of field, and overall correlation were close to the ones observed experimentally (Fig. S11)

#### Position-modulated inputs

The simulated CA3 neurons received feedforward input from *N*^*input*^ neurons whose activities are tuned to specific locations. The location of the peak of the activity for each input neuron was determined by dividing the length of the annular track by the number of input neurons and assigning each neuron to a multiple of this division. Therefore, the firing rate of input neuron *i* was centered at 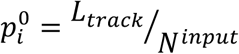, where *L*_*track*_ is the length of the annular track. All the place fields of input neurons had the same tuning width, *σ*_*input*_, and the same amplitude, *A*_*input*_. The animal explored an annular track with speed *v*. The potential variable of input neuron *j* with place field centered at 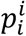 when the animal is at position *p* is

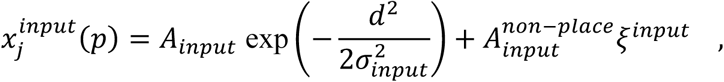

where *d* is the distance, along the track, between the animal’s position and the center of the neuron’s place field, *ξ*^*input*^ is a noise term generated from an Ornstein-Uhlenbeck process with time constant 10 ms, and 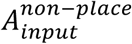 is the amplitude of non-place inputs on input neurons. The value for 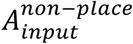 was chosen to fit our experimental data such that the mean within field, outside of field, and overall correlation were close to the ones observed experimentally (Fig. S11).

#### Data-constrained parameters

In all of our simulations, neurons were composed of 100 dendrites directly connected to the soma. The proportion of those dendrites that displayed spatially tuned activity, *p*_*d*_, was extracted from our experimental data and imposed as a constrain to our simulations. These values were extracted from four distinct subsets of our data: tuned cells in familiar environments (*p*_*d*_ = 0.5), untuned cells in familiar environments (*p*_*d*_ = 0.3), tuned cells in novel environments (*p*_*d*_ = 0.17), and untuned cells in novel environments (*p*_*d*_ = 0.06). For tuned cells, we also constrained our model by imposing the proportion of tuned dendrites that were tuned to the same place as the soma (0.65 for familiar and 0.62 for novel environments).

#### Tuned versus untuned cells

In addition to differences in proportions of tuned dendrites, tuned cells were different from untuned cells in terms of efficiency of dendrite-to-soma propagations and distribution of excitatory synaptic weights. For tuned cells in familiar environments, the propagation efficiencies were set to 0.01 for all dendrites and multiplied by a factor of 10 for the dendrites that shared the same tuning as the soma. For untuned cells in both familiar and novel environments, propagation efficiencies were all set to 0.04. Tuned cells in novel environments were assumed to be less tuned than in familiar environment and, therefore, propagation efficiencies were set to 0.04 and multiplied by 3. To define the synaptic weights for tuned cells, we first set all the weights to zero and added a rectified random number drawn from a normal distribution with mean zero (5, for novel environments) and standard deviation 2.0. For each tuned dendrite, the preferred input was randomly assigned (but constrained to the proportion of dendrites tuned to the same location as the soma) and the corresponding synaptic weight was increased by 1000 (150, for novel environments). The weights for each dendrite were then normalized by dividing all the weights by the average weight and multiplying the result by 0.1. For untuned cells, we follow the same steps but the preferred synaptic weight for tuned dendrites is multiplied by 150.

#### Place field stability analysis

We simulated CA3 neurons under two conditions: branch selective tuning—in which all the dendrites were tuned to a random location—and branch prevalence tuning—in which all the dendrites were tuned to the same location. The propagation efficiency for one of the dendrites was multiplied by 10 and this dendrite defined the position of the peak of the somatic place field. The propagation efficiency for all the other dendrites was set to 0.5 for the branch selective simulations, and to 0.1 for the branch prevalence simulations. To enhance the effect of noise, we increased the local noise to dendrites by multiplying the noise amplitude by 2 and reduced all the other sources of noise to zero. The relation between the stability under both conditions considered here—and therefore our conclusions from our simulations—are not dependent on specific values of noise amplitude. To measure the stability of place fields, we first measured the average place field over all trials for each condition considered (50 trials in total for each condition). Place field stability was then defined as the correlation between each trial and the average place field.

#### Place field flexibility analysis

To measure place field flexibility, we simulated CA3 neurons while an animal ran through an annular track for two laps. During the first lap of exploration, the amplitude of the activity of all input neurons was set to the same level (*A*_*input*_ = 0.2). During the second lap, however, one of the input neurons was randomly chosen and its activity amplitude was multiplied by 10 to try to shift the peak of somatic activity to a new location. We simulated CA3 neurons under two conditions: branch selective tuning—in which all the dendrites were tuned to a random location—and branch prevalence tuning—in which all the dendrites were tuned to the same location. Noise amplitudes were set to zero except for local dendritic noise amplitude, which was set to 0.4. The relation between the flexibility under both conditions considered here—and therefore our conclusions from our simulations—are not dependent on specific values of noise amplitude. Place field flexibility was measured as the distance between the position of the initial peak of somatic place field (first lap) and of its final position (second lap) following the increase in amplitude of a randomly selected input place field.

#### Parameters and simulations

All simulations were implemented in python and are available at ModelDB. The parameters used in our simulations can be found in table 2.

**Table S2.** Parameters for simulations.

| **Neuron Model** |  |  |
| --- | --- | --- |
| **Neuron and Network parameters** |  |  |
| **Name** | **Value** | **Description** |
| $N^{dend}$ | 100 | Number of dendritic compartments for CA3 neurons |
| $N^{input}$ | 100 | Number of input neurons |
| $I^d$ | 12 a.u. | Dendritic inhibition |
| $I^s$ | 6 a.u. | Somatic inhibition |
| **Place-tuned input** |  |  |
| **Name** | **Value** | **Description** |
| $A_{input}$ | 2.0 | Presynaptic place field amplitude |
| $\sigma_{input}$ | 2.0 | Presynaptic place field width |
| **Non-place input** |  |  |
| **Name** | **Value** | **Description** |
| $A_{local}^{non-place}$ | 0.4 | Amplitude of local term of non-place inputs |
| $A_{global}^{non-place}$ | 0.4 | Amplitude of global term of non-place inputs |
| $A_{input}^{non-place}$ | 0.2 | Amplitude of non-place inputs onto input neurons |
| **Simulation parameters** |  |  |
| **Name** | **Value** | **Description** |
| $L_{track}$ | 50 a.u. | Track length |
| $v$ | $5.0 \times 10^{-3} \text{ ms}^{-1}$ | Animal speed |

## Author contributions

SKR and JB conceived the project, designed the experiments and wrote the manuscript. SKR performed all mouse experiments and analysis. VP and CC conceived the CA3 model, and VP created the model and all resulting modeling data in addition to experimental analysis in Fig 2I. MAD developed the cranial window preparation and assisted with data analysis. JJM performed analysis for Fig S2-3. RGD performed the immunohistochemistry and confocal imaging for Fig S1. PP and SC conceived and SC implemented the machine learning algorithms for place cell predictions.

## Data and materials availability

All experimental data, modeling code, and machine learning code is available upon request.

