## Supplementary Figures for "The dendritic spatial code: branch-specific place tuning and its experience-dependent decoupling"

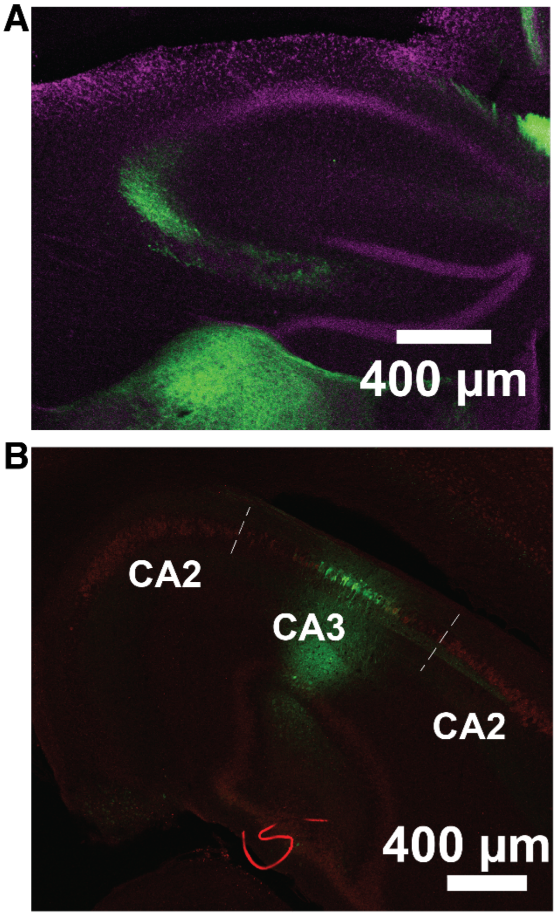


**Fig. S1. Slices from imaged animals.**

**A.** Zoomed in view of hippocampus from Fig 1. GCaMP6f injection into CA3 shown in green. DAPI in magenta. **B.** Horizontal slice from an experimental animal showing GCaMP6f in green and CA2 marker PCP4 in red, non-overlapping. At this level CA2 splits to flank both sides of the ventricle with CA3 in the middle.

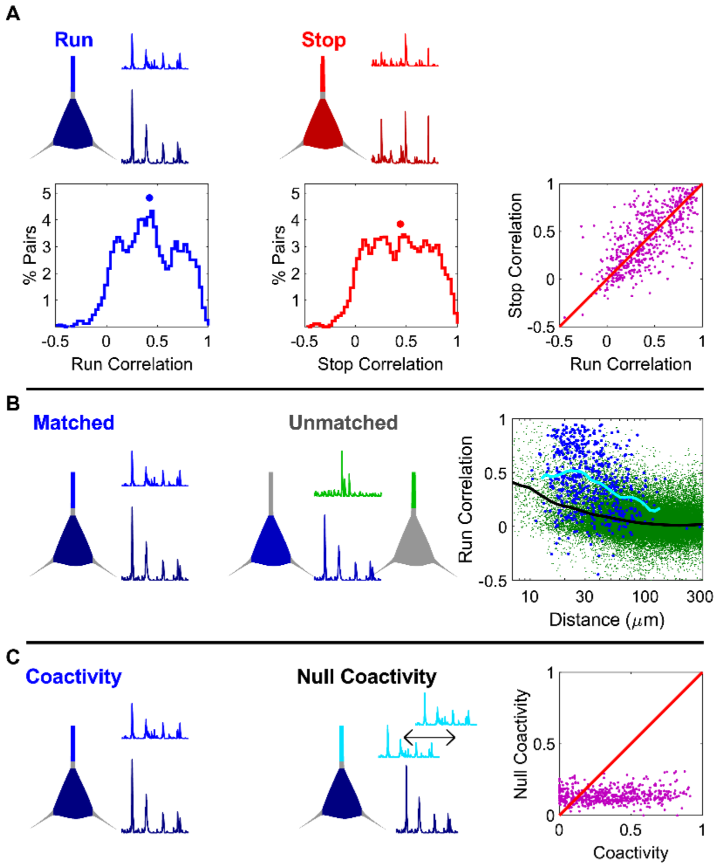


**Fig. S2. Connected soma-dendrite pairs are more correlated than expected by chance.**

a) Top-left, sample traces simultaneously recorded from a soma and dendrite identified as belonging to the same neuron while a mouse is running on the treadmill. Top-right, sample traces from the same soma-dendrite pair during immobility. Bottom-left, histogram of soma-dendrite correlations during running, with a median of 0.42, [0.38, 0.45], n=536 pairs. Bottom-middle, histogram of soma-dendrite correlations during immobility, with a median of 0.44, [0.39, 0.48]. Bottom-right, soma-dendrite correlations during running and immobility were not significantly different (p=0.90, Wilcoxon sign-rank test, n=536 pairs), but they were significantly correlated (r=0.71, p=5.7x10-84, two-sided t test), indicating that differences in correlations are not likely driven by motion artifacts. Median values are expressed as M, [L , U], where M is the median and L and U are the lower and upper bounds, respectively, of the 95% confidence interval of the median obtained through resampling.

b) Left, sample traces as in a top-left. Middle, sample traces from a soma and dendrite belonging to two separate neurons, showing a low degree of correlation. Right, correlation coefficients between soma-dendrite pairs plotted as a function of distance between the ROIs (see methods), for matched soma-dendrite pairs (blue, n=536 pairs) and unmatched pairs (green, n=126017 pairs). The moving average for matched (light blue) and unmatched (black) pairs are overlaid in thick lines. Matched pairs had significantly higher correlations than unmatched pairs (p=3.45x10-65 for group effect, two-way ANOVA with log10(distance) as a continuous predictor).

c) Left, sample traces as in a top-left. Middle, sample traces from the same soma-dendrite pair demonstrating the shifting method used to compute null coactivity (see methods). Right, coactivity of un-shifted soma-dendrite pairs was greater than that of shifted pairs (p=1.2x10-55, Wilcoxon sign-rank test, n=536 pairs). The un-shifted coactivity was greater than the shifted coactivity for 79% (441/559) pairs.

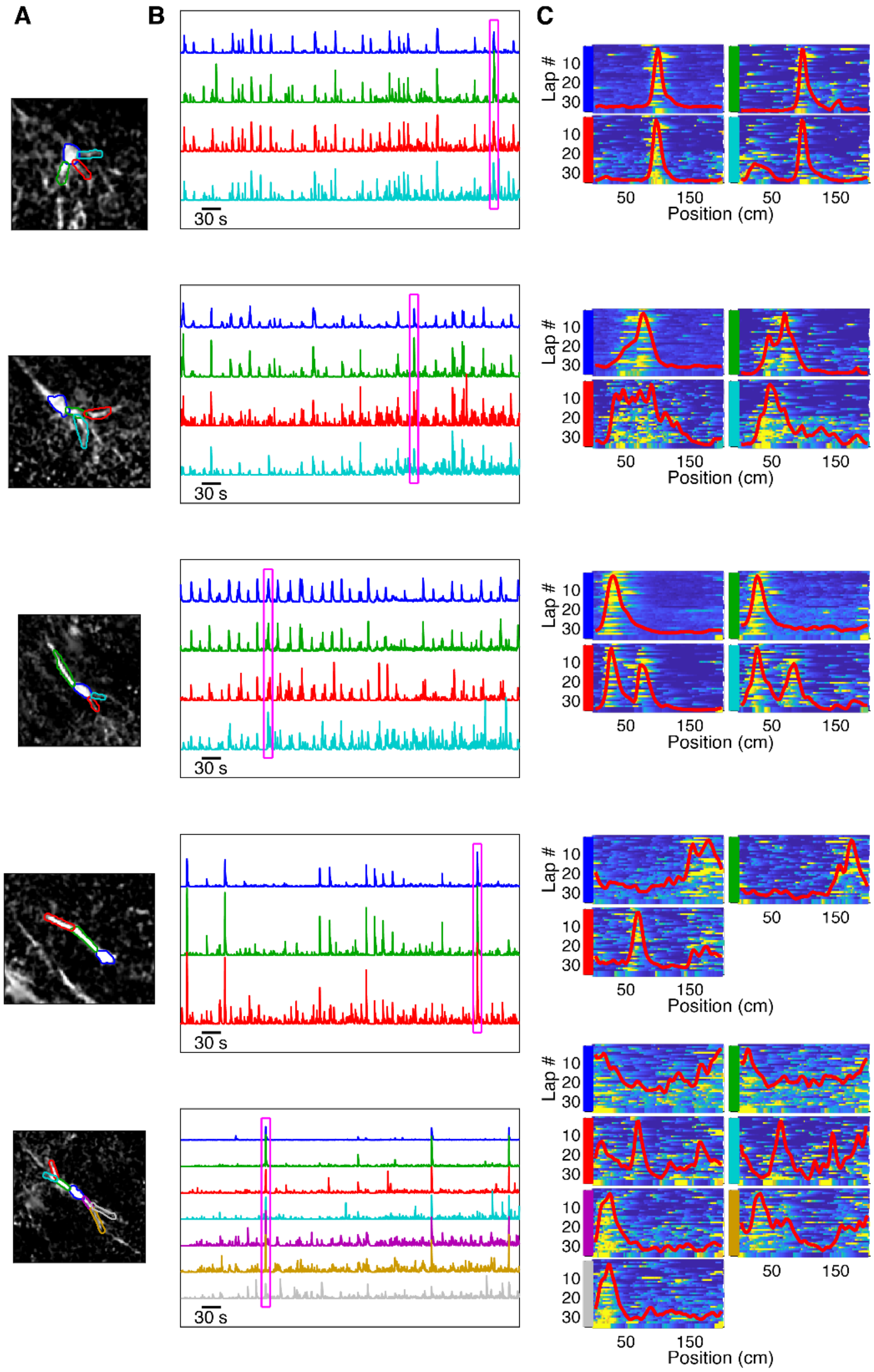


**Fig S3. Examples of simultaneous events across multiple compartments.**

a) Spatial footprint of soma and connected dendritic segment ROIs. Note that all dendritic ROIs are contiguous in space with their soma, indicating all ROIs belong to the same neuron. b) Sample raw traces of the ROIs identified in a. The magenta box indicates synchronous activity, with high ΔF/F in all compartments. Note all example cells exhibit multiple globally synchronous events, indicating all ROIs belong to the same neuron. c) ΔF/F as a function of lap number and position for all ROIs identified in a. The tuning curve averaged over all trials (see methods) is overlaid in red. Note that several dendrites, particularly those in the bottom two examples, show substantially different tuning than their parent soma.

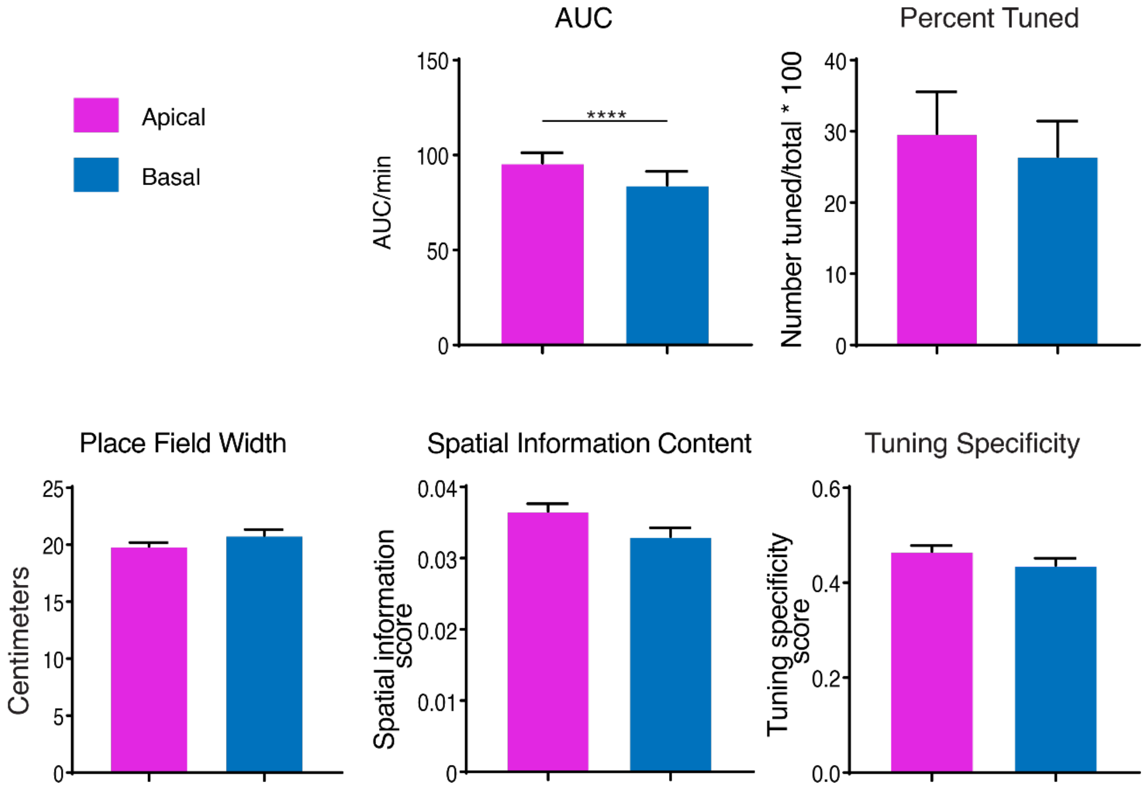


**Fig. S4. Apical dendrites are slightly more active, but are otherwise no different from basal dendrites in terms of spatial tuning.** Apical dendrites are more active than basal dendrites (median apical AUC = 63.97, median basal AUC = 49.03, $p < 0.0001$, Mann-Whitney test). The percent tuned (median apical percent tuned = 38.71%, basal percent tuned = 24.00%, $p = 0.5776$, Mann-Whitney test), place field width (median apical width = 18.27 cm, median basal width = 18.70 cm, $p = 0.2690$, Mann-Whitney test), spatial information score (median apical spatial information = 0.03121, median basal spatial information = 0.02826, $p = 0.0519$, Mann- Whitney test), and tuning specificity score (median apical tuning specificity = 0.4304, median basal tuning specificity = 0.3957, $p = 0.1517$, Mann-Whitney test) are no different.

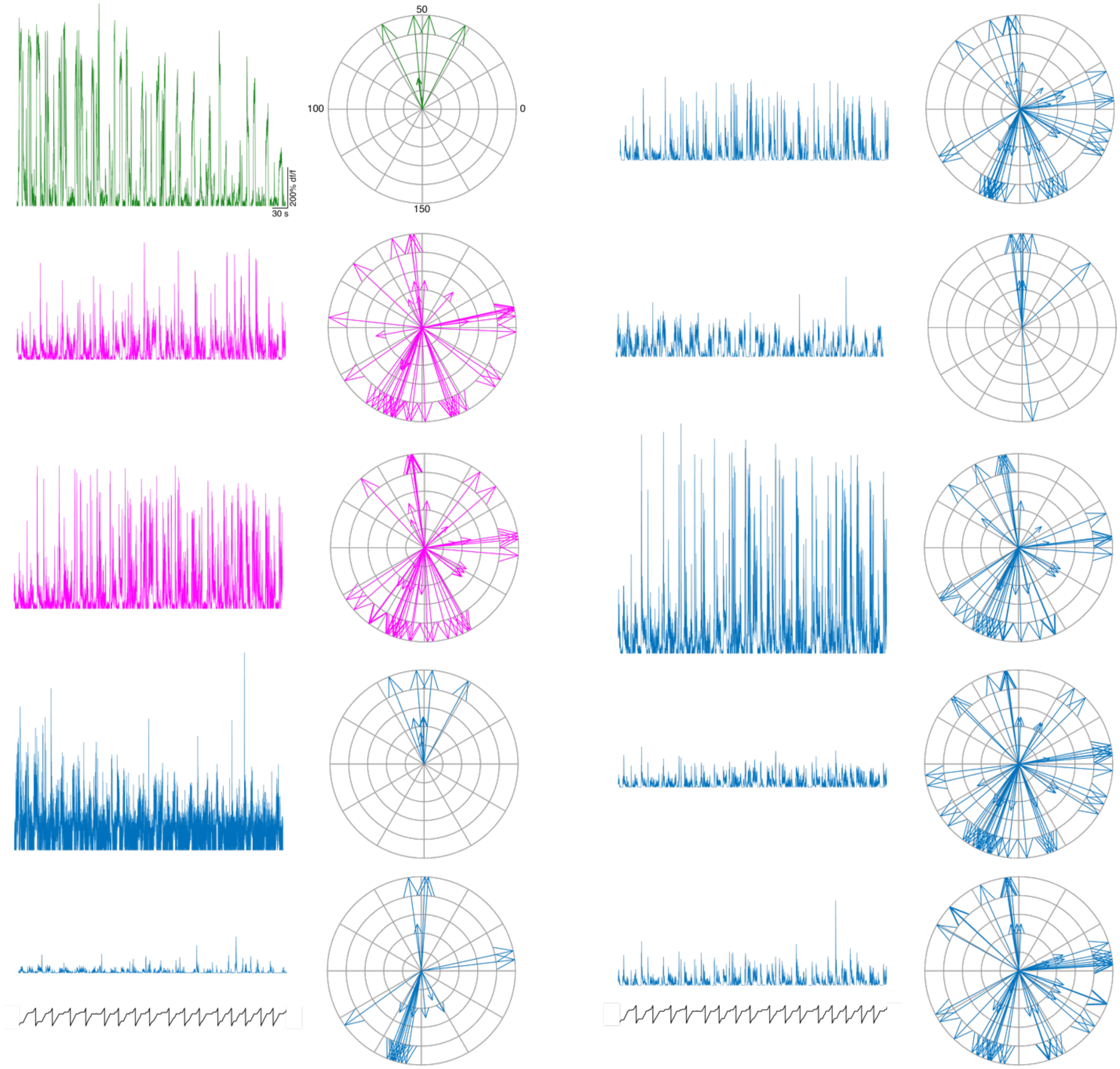


**Fig. S5. Sparsely expressing dataset shows branch-specific tuning.** df/f traces and tuning vector plots for cell shown in 1H.

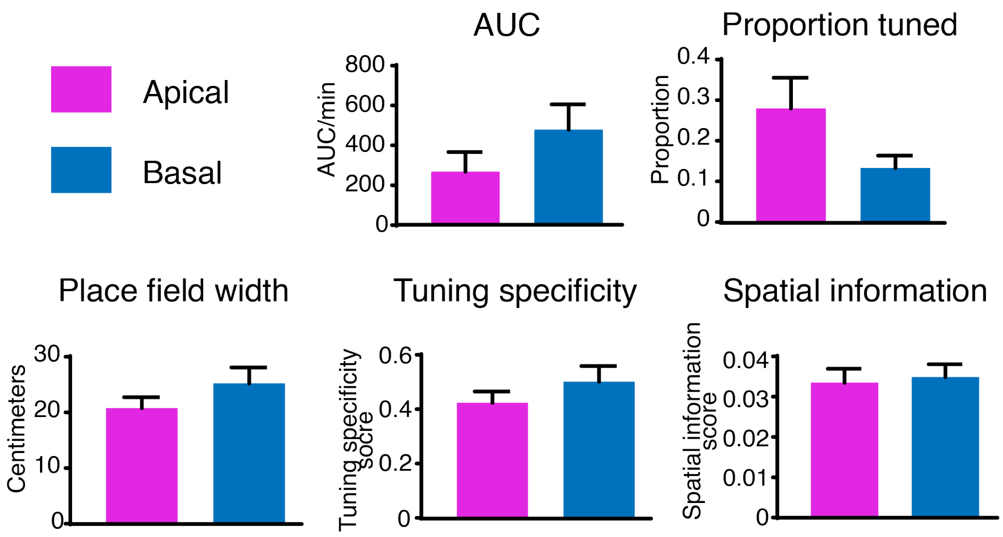


**Fig. S6. No differences observed between tuning properties of apical and basal dendrites in a sparse GCaMP expressing dataset.** Similar to our findings in our densely expressed GCaMP dataset, we found no significant differences in activity rate (basal = 111.2, apical = 210.5, $p = 0.13$; Mann-Whitney test), proportion tuned (basal = 0.1622, apical = 0.2440, $p = 0.1265$; Mann-Whitney test), place field width (basal = 20.50 cm, apical = 18.60 cm, $p = 0.3940$; Mann-Whitney test), tuning specificity score (basal = 0.4710, apical = 0.4411, $p = 0.4643$; Mann-Whitney test), or spatial information score (basal = 0.03273, apical = 0.03356, $p = 0.9285$; Mann-Whitney test).

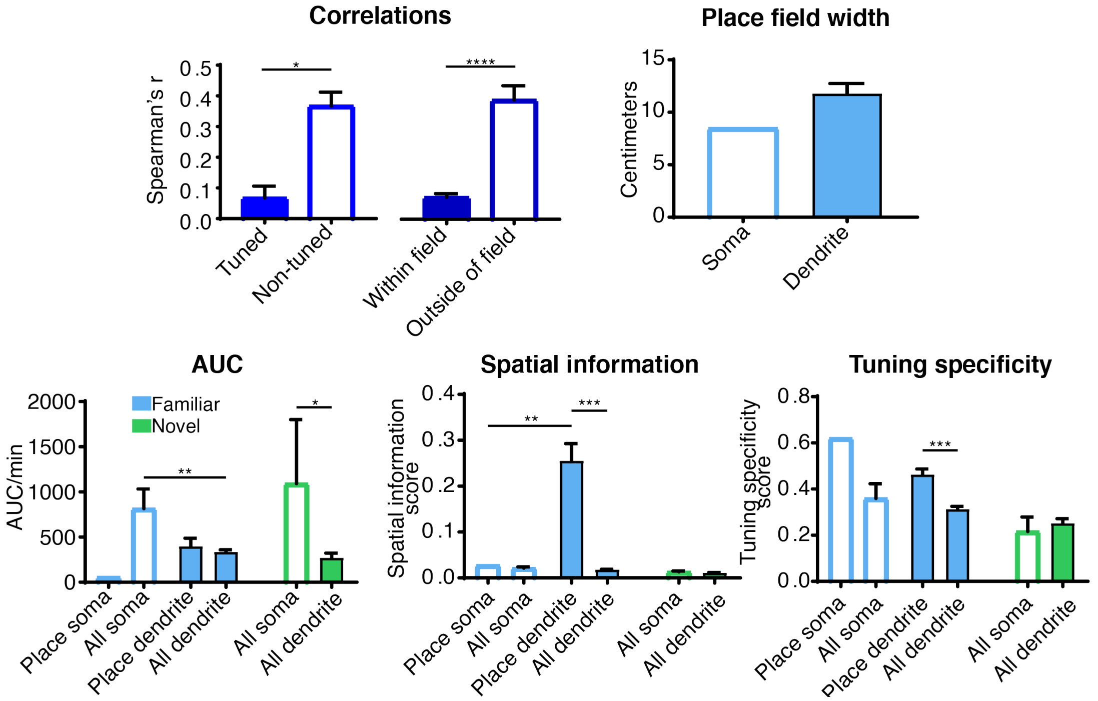


**Fig. S7. Tuning metrics and correlation relationships are similar between dense and sparse GCaMP expressing dataset.** We saw the same soma-dendrite correlation relationships in the familiar environment in our sparsely-expressing dataset as our densely-expressing dataset (tuned Spearman's r = 0.03691, non-tuned = 0.4545, p = 0.0374, Mann-Whitney test; outside of field = 0.06706, in field = 0.4626, p < 0.0001, Mann-Whitney test). We did not have any tuned compartments in the novel condition, but we saw similar values in the familiar condition: place field width (soma = 8.46 cm, dendrite = 11.77 cm), activity rates (familiar place soma = 57.32, familiar place dendrite = 399.4, p = 0.9913; familiar all soma = 816.5; familiar all dendrite = 335.5; familiar place soma vs all soma, p = 0.6373; familiar place dendrite vs all dendrite, p = 0.9848; familiar all soma vs all dendrite, p = 0.0015; novel all soma = 1093, novel all dendrite = 271, p = 0.0317; familiar all dendrite vs novel all dendrite, p = 0.9124; familiar all soma vs novel all soma, p = 0.9619; one-way ANOVA with Sidak's multiple comparisons), spatial information (familiar place soma = 0.02797, familiar place dendrite = 0.2555, p = 0.0016; familiar all soma = 0.02154, familiar all dendrite = 0.01827; familiar place soma vs familiar all soma p>0.9999; familiar place dendrite vs familiar all dendrite p<0.0001; novel all soma = 0.01351, novel all dendrite = 0.01113; familiar all dendrite vs novel all dendrite p = 0.9221; familiar all soma vs novel all soma, p =0.9913; novel all soma vs novel all dendrite, p > 0.9999; one-way ANOVA with Sidak's multiple comparisons), and tuning specificity (familiar place soma = 0.6213, familiar place dendrite = 0.4635, p = 0.9789; familiar all soma = 0.3594, familiar all dendrite = 0.3128; familiar place soma vs familiar all soma, p = 0.8131; familiar place dendrite vs familiar all dendrite, p = 0.0004; novel all dendrite = 0.2521, novel all soma = 0.2162, p = 0.9999; familiar all dendrite vs novel all dendrite, p = 0.1470; familiar all soma vs novel all soma p = 0.8805; one-way ANOVA with Sidak's multiple comparisons).

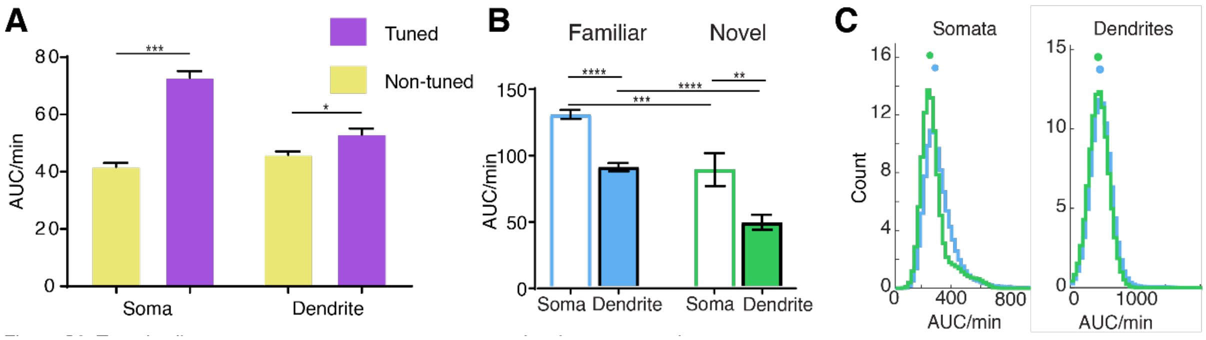


**Fig. S8. Tuned cell compartments are on average more active than non-tuned compartments.**

**A.** For both somata and dendrites, activity of tuned cell compartments (mean tuned somatic AUC = 72.76, mean tuned dendritic AUC = 52.97) is higher than activity of non-tuned compartments (mean non-tuned somatic AUC = 41.73, mean non-tuned dendritic AUC = 45.90) ($p < 0.0001$ for soma, $p = 0.0161$ for dendrites , One way ANOVA). This likely results from some cells being silent (and therefore non-tuned). **B.** Both place somata and place dendrites have larger calcium transients in the familiar environment ($p<0.001$ for somata, $p<0.0001$ for dendrites, Mann Whitney test with Bonferroni correction for multiple comparisons) suggesting that spatially tuned activity is more strongly driven in the familiar environment as compared to the novel (Fig S4). In any environment, place somata have greater mean fluorescence than place dendrites (mean familiar somatic AUC/min = 126.7, mean familiar dendritic AUC/min = 87.51, $p<0.0001$; mean novel somatic AUC/min = 86.04, mean novel dendritic AUC/min = 46.83, $p<0.01$ Mann-Whitney test with Bonferroni correction for multiple comparisons), perhaps an outcome of burst firing that is characteristic of place somata (14,25, 32). **C.** Histogram showing distributions of AUC for place somata and dendrites in familiar vs. novel environments.

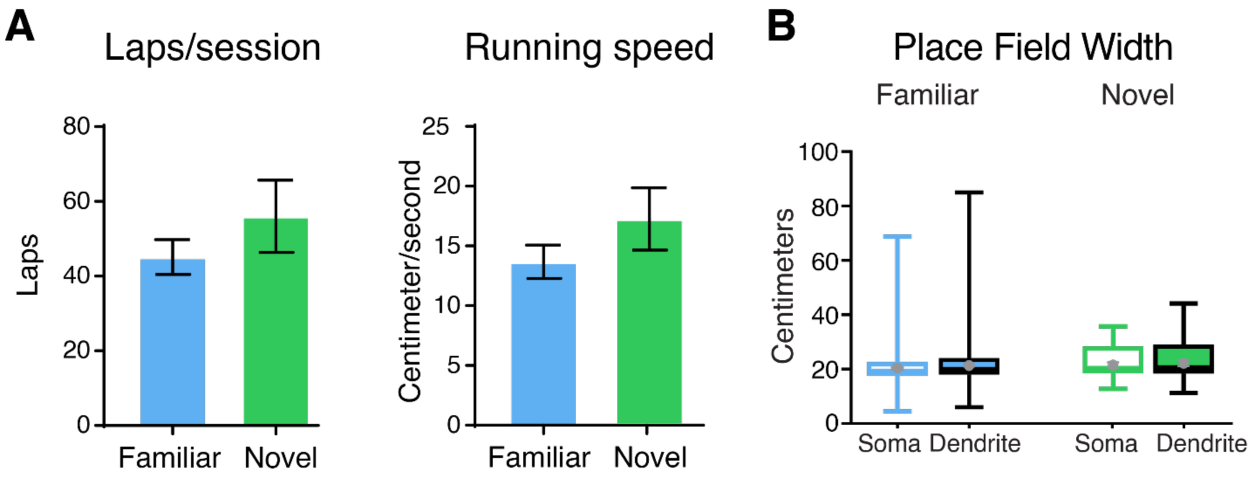


**Fig. S9. No differences in number of laps run, running speed, or place field width between familiar and novel environments.** Difference in novel and familiar spatial coding were not due to differences in number of laps run (familiar = 45.07 laps/session, novel = 56 laps/session, $p = 0.7319$; Mann-Whitney test) or speed of the animal (familiar = 13.67 cm/s, novel = 17.11 cm/s, $p = 0.7011$; Mann-Whitney test). Additionally, place field widths are not different between somata and dendrites both within and across conditions.

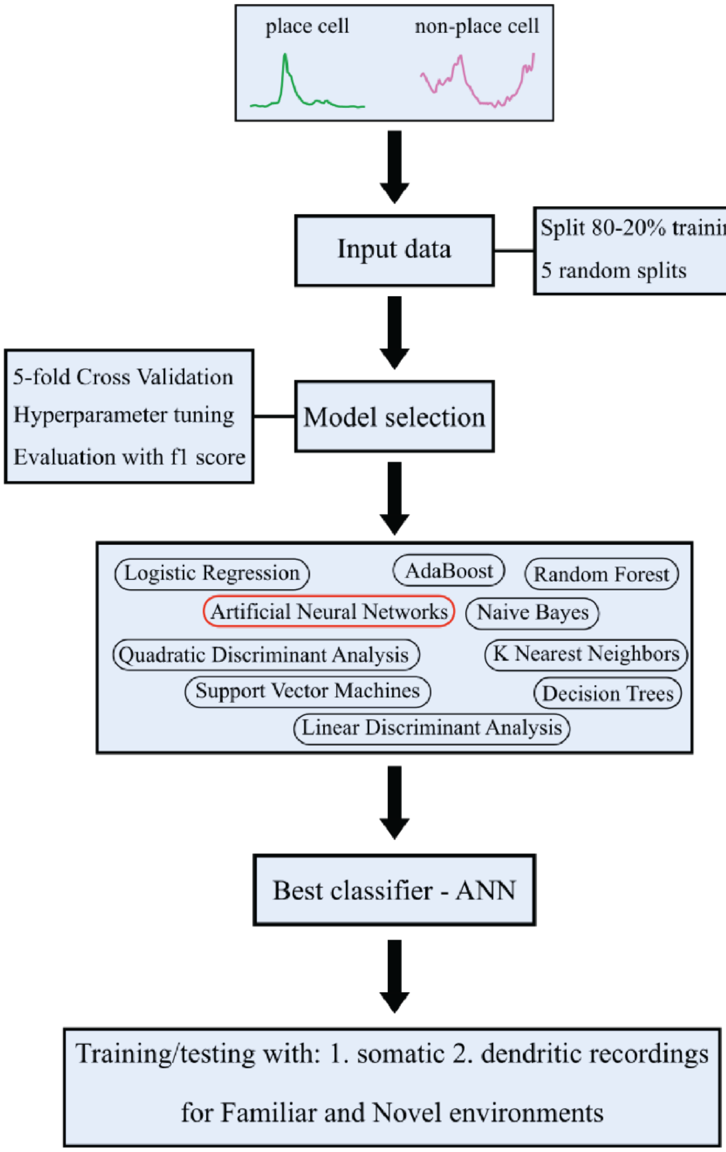


**Fig. S10. Schematic diagram of the Classification process.** We used somatic recordings from both place (green) and non-place (magenta) cells in the familiar environment as input to various classifiers, in order to select the best classifier and find its optimal hyperparameters. To do this, we split the dataset into training (80%) and test (20%) sets, respectively. We then used 5-fold Cross Validation to find the best hyperparameter set for each classifier (the 5-fold CV was repeated for all hyper-parameter combinations, i.e., grid search). Specifically, the process was repeated 5 times (5 random splits) and the classifier with the highest average f1 score was selected. The selected algorithm (with the best set of hyperparameters) was used to classify the cells into place and non-pace cells using four datasets, i.e., somatic recordings in the familiar environment, somatic recordings in the novel environment and dendritic recordings in familiar and novel environments, respectively. The datasets were again randomly split into training (80%) and test (20%) sets, respectively and the algorithm was trained and tested. The process was repeated 100 times and the mean AUC score was calculated on the test set. To ensure balanced classes, we downsampled each time the data set by removing data from the majority class (non place cells) in order to have 50-50 class representation.

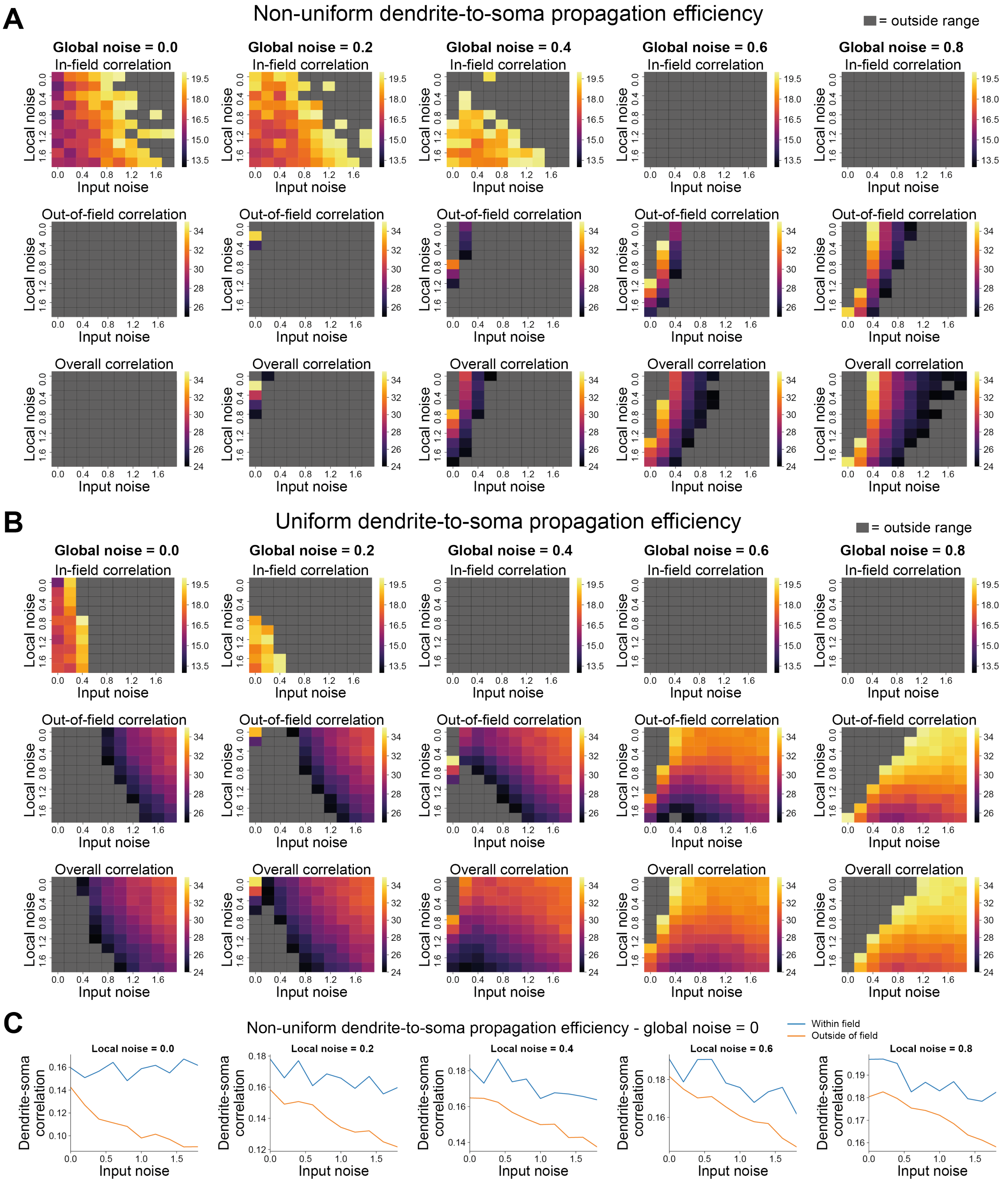


**Fig. S11. Experimental data-fitting in simulations suggests non-uniform dendrite-to-soma propagation efficiency. A.** Within field correlation (top row), outside of field correlation (middle row), and over all correlation (bottom row) between dendrites and soma for tuned cells in familiar environments (all the values are multiplied by 100). Each value shows an average over 50 trials of a simulated CA3 neuron with non-uniform dendrite-to-soma propagation efficiency. Each column shows the results for one level of global noise (see methods). Within field correlations were considered when within the range from 13 to 20. Outside of field correlations were considered when within the range from 25 to 35. Overall correlations were considered when within the range from 24 to 35. Grey boxes indicate values outside of these ranges. **B.** Same as in A but for simulations of CA3 neurons with uniform dendrite-to-soma propagation efficiency. We could not find any values of local, global, and input noise that would simultaneously satisfy the constraints imposed by our experimental data. Therefore, these simulations suggest that the efficiency of dendrite-to-soma propagation is not homogeneous across all the dendrites. **C.** Within field (blue) and outside of field (orange) dendrite-soma Spearman's correlation as a function of input noise (non-place tuned input at input neurons) for simulations of CA3 neurons with non-uniform dendrite-to-soma propagation efficiency and global noise (global non-place tuned input) set to zero. Each column is related to one level of local noise (local non-place tuned input). For all levels of local and input noise, within field correlation is higher than outside of field correlation if the level of global noise is set to zero.

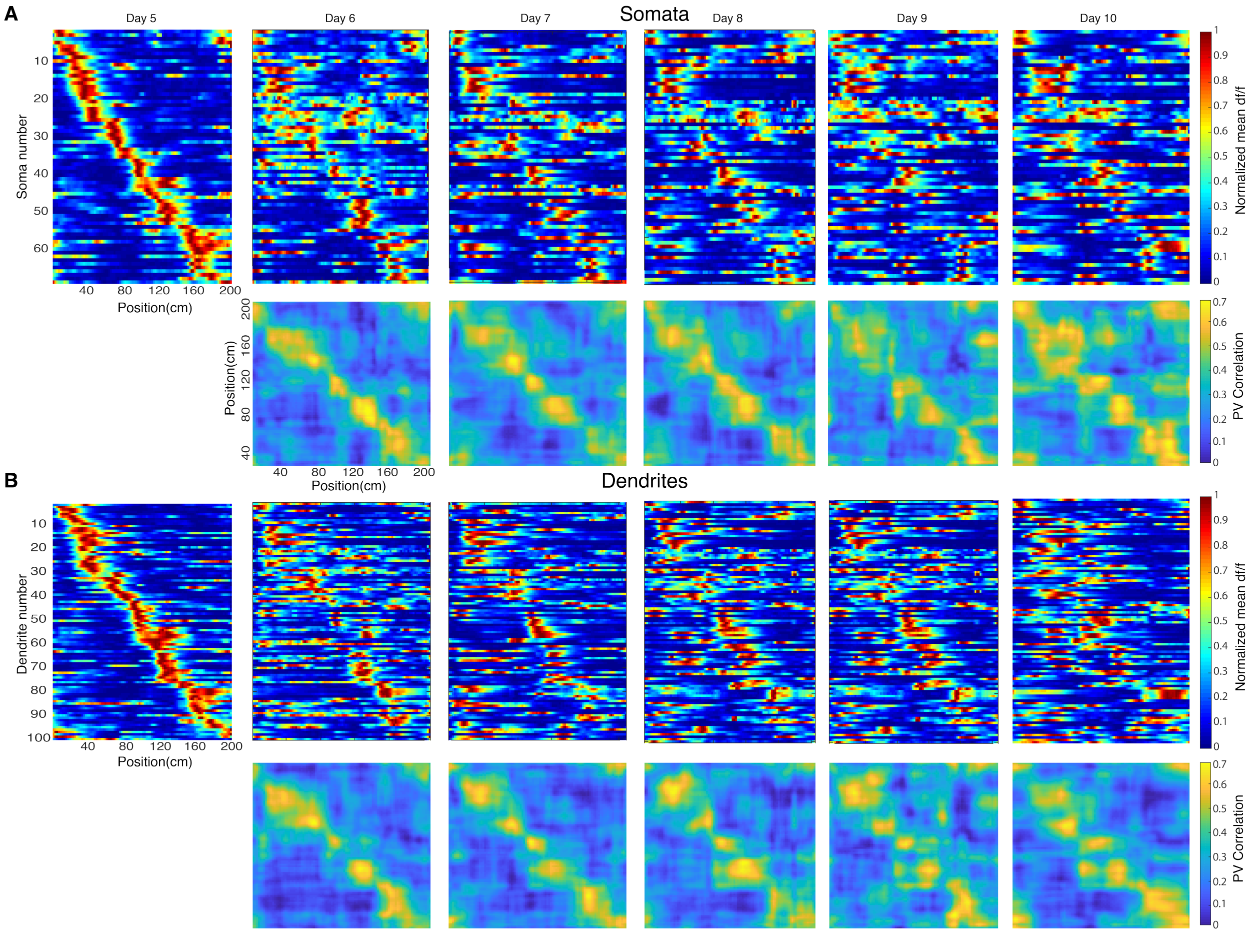


**Fig. S12. Place dendrites and somata show no difference in ensemble stability across time.** a) Top: Example normalized mean df/f for tuned somata on day 5, and those same somata through day 10. Bottom: Population vector correlations for days 6-10 as compared to day 5. b) Top: Example normalized mean df/f for tuned dendrites on day 5, and those same dendrites through day 10. Bottom: Population vector correlations for days 6-10 as compared to day 5.

**Movie S1.**
Example imaging video from our dense dataset temporally downsampled by a factor of 20 for presentation purposes only.

**Movie S2.**
Example imaging video from our sparse dataset temporally downsampled by a factor of 20 for presentation purposes only.
